# Jumping spiders are not fooled by the peripheral drift illusion

**DOI:** 10.1101/2024.09.17.613447

**Authors:** Massimo De Agrò, Giorgio Vallortigara, Egidio Falotico

## Abstract

In the peripheral drift illusion, a static circular sawtooth pattern is perceived as if it were rotating. It is believed that this effect is a byproduct of how the neural substrate responsible for motion perception is organized. The structure of the motion perception circuitry is widespread across the animal kingdom, vertebrates and invertebrates alike, which in turn causes the illusion effect to be experienced by virtually all animals. Among invertebrates, jumping spiders possess a unique visual system. For them, the tasks of visual computation are split across 4 pairs of eyes, with motion detection, target recognition, and shape discrimination computed in completely segregated brain areas and visual field sections. In such an organization, it is unlikely that the circuitry for motion perception common to other animals is shared by jumping spiders. Consequently, jumping spiders should be immune to the peripheral drift illusion. To test this hypothesis, we placed jumping spiders on top of an omnidirectional treadmill and presented them with circular visual stimuli in their visual periphery. These were either composed of a sawtooth pattern, and therefore inducing the illusion, or of a sine-wave pattern of equal luminance and spatial frequency but not illusion-inducing. The stimuli could either be static or rotate around their center, either clockwise or counterclockwise. As jumping spiders perform distinctive full-body pivots when detecting a moving object in their visual periphery, we registered the frequency of this behavior to assess the illusory percept. We found that the spiders responded consistently to all moving stimuli, but did not react to the static illusion, therefore it was not perceived as in motion. The absence of the illusory percept in spiders opens many questions about the nature of their motion perception circuitry and casts doubts on how the illusion is widespread in the animal kingdom outside the common model species usually inquired about.

## Introduction

In the struggle for survival, animals have developed a wide variety of sensory system, in order to rapidly and efficiently interpret their surrounding environment. One of the most widely employed and effective of these senses is vision. Thanks to photo-sensitive cells capable of detecting incoming photons, our brain can extract information about the objects that surround us, their shape, and their movement (Lazareva et al., 2012). However, no visual system can provide a perfectly accurate representation of the quick enough to make relevant decisions. Instead, brains take computational shortcuts, using simple rules to infer what is happening in the environment, without needing perfect information.

These shortcuts are most evident when errors in perception occur. Some visual stimuli, commonly referred to as “visual illusions”, exploit the inner-working of the visual system, generating the perception of a certain characteristic even if this is physically absent (Todorović, 2020). These illusions have been the focus of scientific inquiry in experimental psychology for the last century (Kanizsa, 1979), as the nature of the perceptual error can reveal the mechanisms of visual perception in the brain (Eagleman, 2001). While studies have been mostly focused on humans, more and more experiments have been performed on visual illusions in other animal species, providing great insight into the similarities or differences among phylogenetically distant visual systems. For example, it has been observed that the rules that govern the segregation of objects from the background is shared across mammals, birds, fish, and insects (Kelley & Kelley, 2014; Nieder, 2002; Rosa Salva et al., 2014; Vallortigara, 2004).

While many illusory stimuli provide erroneous percept in the shape of objects, some others pertain to motion. For example, some stimuli can appear to move differently from the physical reality (in terms of speed, direction or others), while, even more strikingly, others can appear to move even if they are in fact completely static. This is the case of the peripheral drift illusion, first described by Fraser and Wilcox (1979), and subsequently by Faubert and Herbert (1999): a luminance sawtooth pattern organized in a circle induces the impression of a rotation in the same direction as the dark to light gradient (Figure 1). Over the decades, many variants of the same illusory stimulus have been developed, employing a step-wise rather than a sawtooth pattern (Kitaoka & Ashida, 2003) or by using color (Kitaoka, 2003). Likewise, the illusion has been extensively studied in humans and other vertebrates (Agrillo et al., 2015; Bååth et al., 2014; Gori et al., 2014).

**Figure 1.**
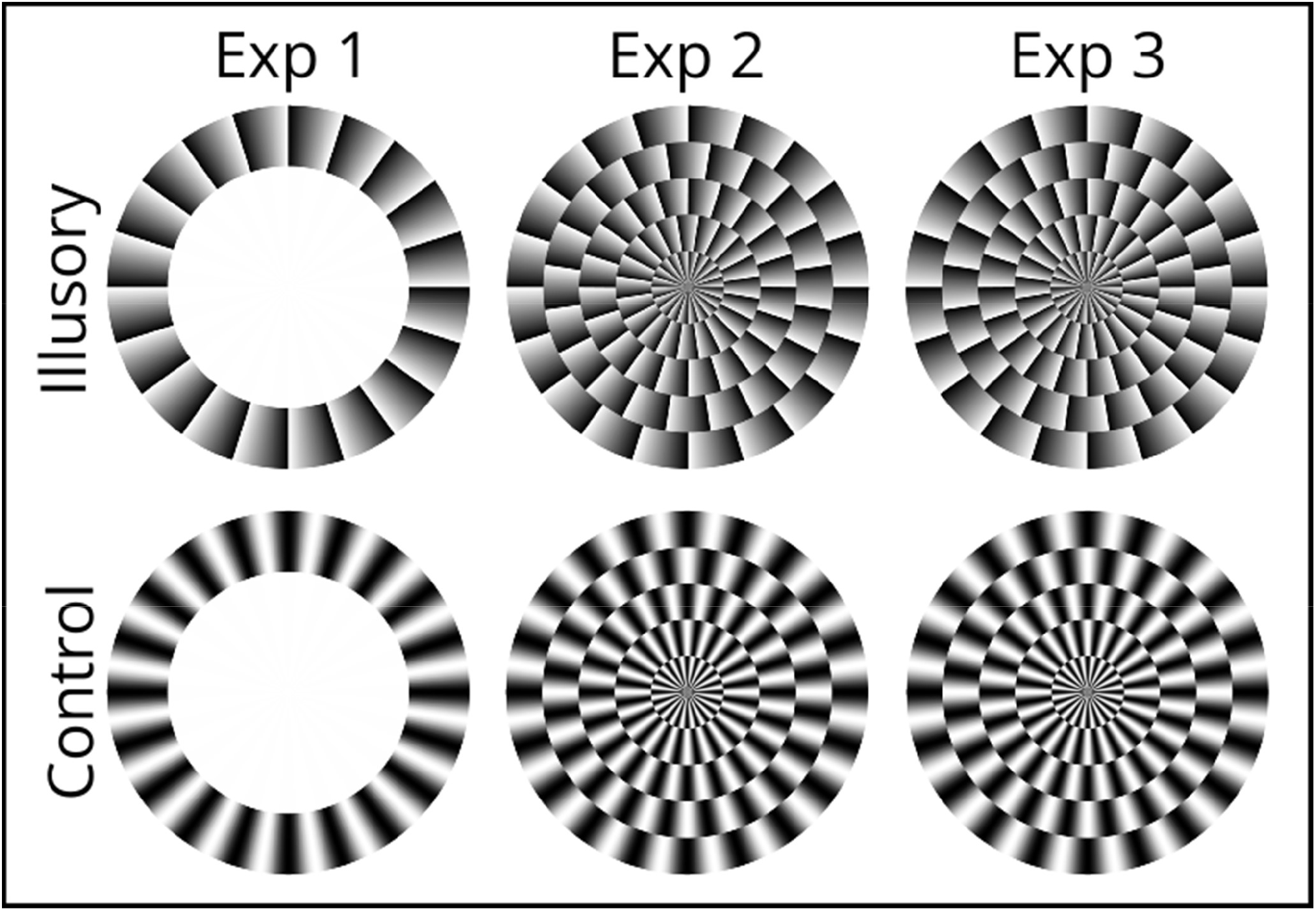
Stimuli used in the three experiments. Illusion-inducing stimuli are characterized by a sawtooth pattern, going from light to dark and then drop. The control stimulus instead presents a sine-wave pattern. In experiment 1, the two stimuli were constituted of a ring pattern, while in experiment 2 and 3 they were made of 5 concentric rings. Experiment 3 had the illusory stimulus flipped, in turn reversing the direction of the illusory rotation. The peripheral drift illusion according to the literature induces a motion in the direction of dark to light (Agrochao et al., 2020; Faubert & Herbert, 1999; Hsieh et al., 2006). This should appear as rotating clockwise in experiment 1 and 2, but counter-clockwise in experiment 3.

**Figure 2.**
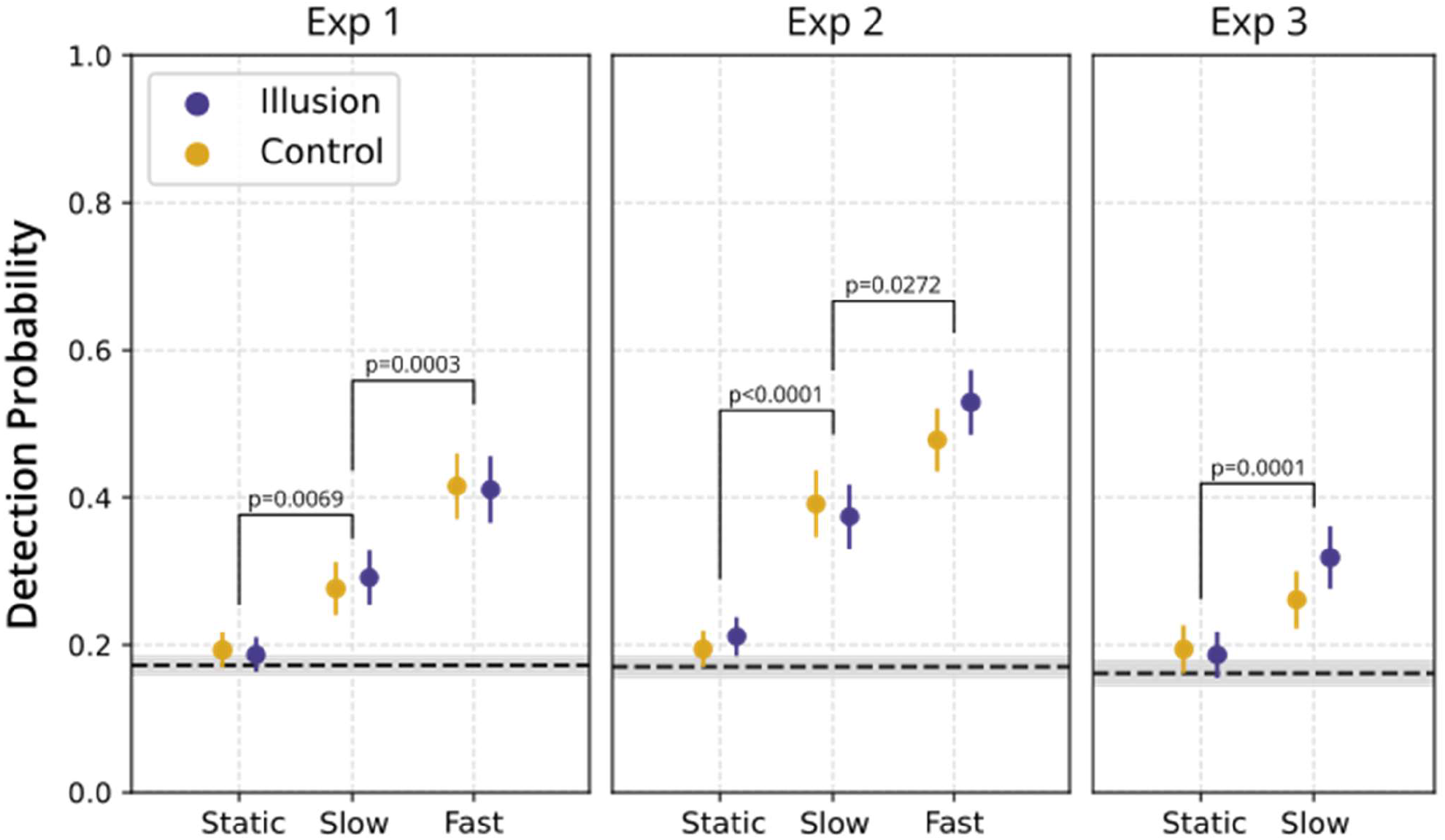
Results. The three plots represent the three experiments. On the X-axis, the different motion speed: (i) static, (ii) 2° per second rotation (slow), (iii) 32° per second rotation (fast). On the Y-axis, the probability of the spider to turn upon stimulus appearance, as calculated by the model. Error bars represent standard errors. The grey dashed horizontal line represents the observed response probability of the spider when no stimulus was presented, considered as a baseline activity. The shaded grey area represents the standard error for this measure. P values are reported for significant contrasts. Across all three experiments, we observed no difference between control and illusory stimuli for none of the tested speed. Moreover, for all three experiment the detection probability of static stimuli did not significantly differ from the responses in case of no stimuli appearance. Across the three experiments we consistently observed an increased detection probability for faster moving stimuli.

**Figure 3.**
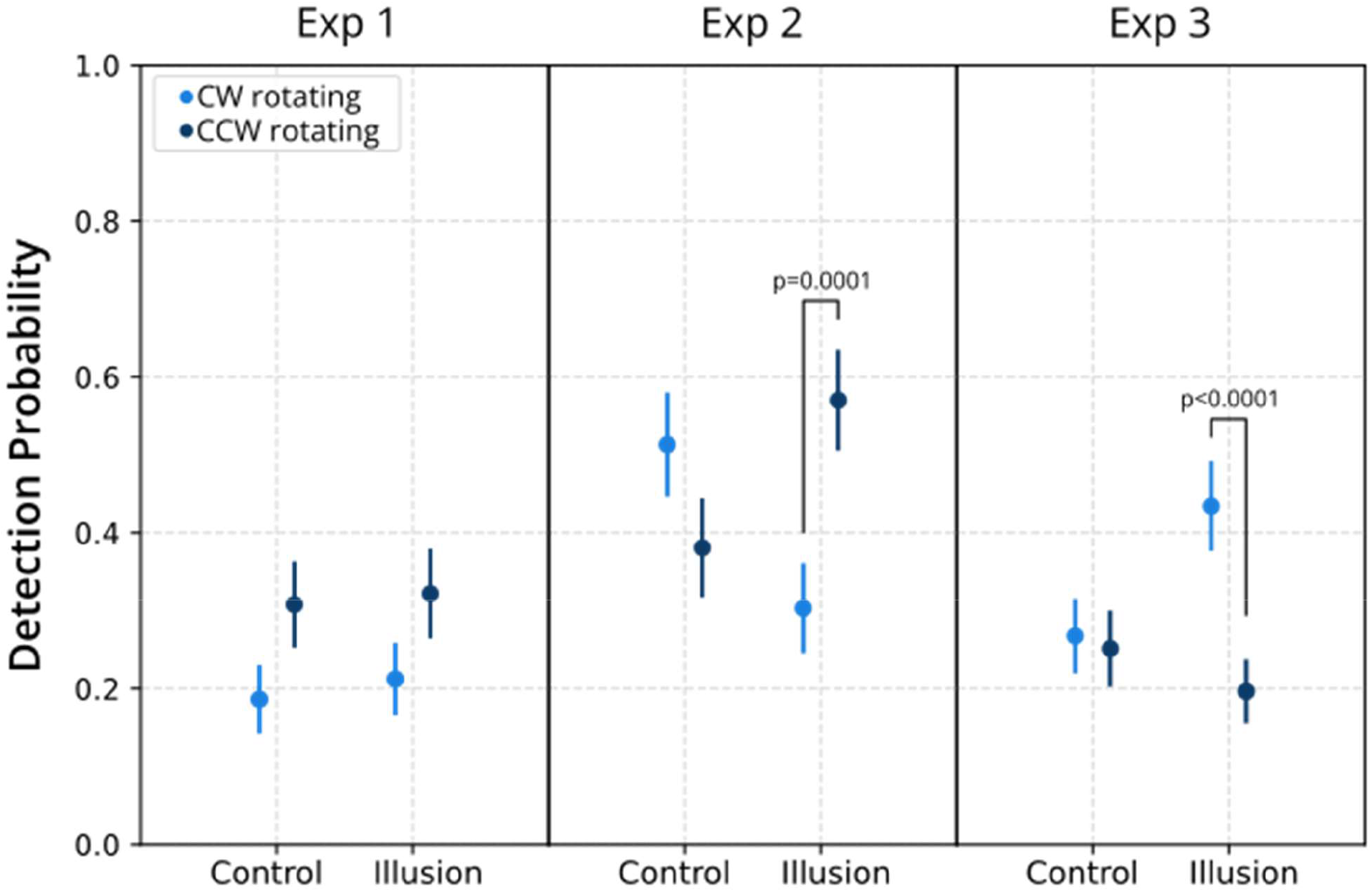
Effect of rotation direction. The three plots represent the three experiments. On the X-axis, we report the different stimuli: either illusory or control. Only stimuli rotating at 2° per second (slow) are shown here, as fast-moving ones showed no effect, and static ones did not have a rotation direction. The different colors represent the different rotation directions, Clockwise (CW) or Counter-Clockwise (CCW). On the Y-axis, we report the probability of the spider to turn upon stimulus appearance, as calculated by the model. Error bars represent standard errors. P values are reported for significant contrasts. In experiment 1, we observed a significant higher detection probability for CCW rotating stimuli across both conditions. This effect however disappears when considering each condition alone, probably due to the halved n. In experiment 2, spiders detected significantly more CCW motion, but only for the illusory stimuli. In experiment 3, where we mirrored the illusory stimuli, the effect reversed, with spiders detecting significantly more CW motion.

Two main, non-mutually exclusive mechanisms have been hypothesized to cause the illusory motion perception in the peripheral drift. The first hypothesis argues that high contrast or high luminance stimuli are processed faster. As such, hard light-to-dark edges of the illusory stimulus appear to be presented before than the neighboring sections, generating an illusion of motion (Conway et al., 2005; Faubert & Herbert, 1999). The second hypothesis attributes instead the illusion to the eye movements that constantly produce small changes in the perceived image, causing the impression of a rotating stimulus (Murakami et al., 2006; Otero-Millan et al., 2012).

More recently, the study of the peripheral drift illusion has been extended to invertebrates. Agrochao et. al. (2020) exploited the optomotor response of *Drosophila* to test whether this Diptera perceive a rotating motion from a sawtooth pattern presented all around its visual field. The authors found that fruit flies do indeed respond to the peripheral drift illusion, and do so in a direction consistent with what has been observed in humans and other vertebrates. Then, by selectively silencing T4 and T5 neurons in the *Drosophila* visual system, the authors successfully demonstrated that the illusory motion perception is caused by an imbalance in the ON and OFF edges motion pathways. Finally, authors discovered evidences supporting the presence of the same mechanism in humans. This introduces a third hypothesis for the perception of motion in the peripheral drift stimulus, which appears to be valid across different taxa.

Over the last decades, a new invertebrate model has fascinated scientists for its unique visual system organization, the jumping spider. These arachnids possess eight different eyes, divided into two separate functional sets (Chong et al., 2024; Winsor et al., 2024). The two anterior medial eyes (AMEs) are usually referred as principal eyes (Strausfeld et al., 1993). They have a high visual acuity, while being characterized by a narrow visual field of only 5° each (Land, 1985). Thanks to a set of muscles however the spider can move the back of the long AMEs eye tubes, effectively shifting their visual fields up to a maximum span of about 60° (Land, 1969). It is widely believed that AMEs are specialized in pattern recognition: when presented with a target, jumping spiders shift their AMEs gaze towards it and engage in a “scanning” behavior, where the AMEs field is moved across the figure, inspecting its shape (Dolev & Nelson, 2014; Land, 1969; Rößler et al., 2022; Winsor et al., 2021). The other three sets of eyes, anterior lateral, posterior medial and posterior lateral (ALE, PME, PLE) are usually referred as secondary eyes (Strausfeld & Barth, 1993). They have a lower acuity compared to AMEs and are unmovable. However, their combined visual field cover almost 360° around the animal (De Agrò et al., 2024; Land, 1985). Secondary eyes are responsible for motion detection: when a moving target is detected across their visual field, the spider promptly rotates its body to face it frontally, preparing for engagement by the AMEs (Bruce et al., 2021; Jakob et al., 2018; Zurek & Nelson, 2012b, 2012a). It has been observed that the secondary eyes are not simple motion detectors (Beydizada et al., 2024; Bruce et al., 2021; Ferrante et al., 2023; Zurek et al., 2010): they can in fact pick and select which stimulus requires further inquiry among multiple options (De Agrò et al., 2021), with each eye-pair (ALE, PME and PLE) specialized in a different motion-based type of discrimination (De Agrò et al., 2024).

No data is available about motion illusion perception in jumping spiders, though the presence or absence of it would provide precious information. First, being the motion-detection eyes all unmovable, the perception of the illusion in stimuli presented to ALE, PME or PLE would conclusively exclude any role of resting eye movements (even though this role is already uncertain, given the recent findings of Agrochao et al., 2020). On the other hand, the perception of the illusion may provide crucial information about the spider’s visual system structure, as it would imply the presence of an ON and OFF edge system, similar to the one observed in *Drosophila*. Conversely, if jumping spiders do not perceive the illusion, this would suggest that in these animals the motion-detection system functions in a fundamentally different way than that observed in vertebrates and invertebrates alike. We hypothesize that indeed jumping spiders will not perceive the peripheral drift stimuli as moving, for given the modularity of their visual system, it is possible that every eye does not need two parallel circuits for light and dark edges, with instead every different eye specialized in either one or the other edge type.

To test the presence of illusory perception in these spiders, we designed three behavioral experiments exploiting the full body rotations that the spider performs when detecting a moving object in the secondary eyes’ fields of view, as described above.

In the first experiment, we presented spiders with two different stimuli (Figure 1). One constituted of a sawtooth luminance pattern arranged as a ring, inducing an illusory clockwise rotation percept. The other stimulus was constructed with a sine-wave rather than a sawtooth pattern, disrupting the illusion while maintaining all low-level features. The stimuli presented could either be static, rotating slowly, or rotating rapidly. For the ones rotating, this could happen either clockwise (i.e., congruently with the illusory percept) or counterclockwise.

In the second experiment, we aimed at augmenting the illusory effect by incrementing the number of light-dark edges. We achieved this by inserting four more rings in the stimulus, shifted by half a phase, effectively constructing concentric versions of the experiment 1 versions (Figure 1).

In the third experiment, we flipped the stimuli of Experiment 2, effectively reversing the illusory percept (which should not induce a counterclockwise rotation).

In all three experiments, for every stimulus presented, we recorded whether the spiders turned to face it or not as a proxy for the detection of a moving target.

## Results

Only the main results are reported here. For the full analysis see SI2.

### Experiment 1 – Annular peripheral drift illusion

In the first experiment, the spiders were presented with annular stimuli, being either static or in motion, and inducing an illusory motion effect or not (control).

The first model revealed an effect of motion speed on the detection probability (GLMM analysis of deviance, χ^2^=80.1041, DF=2, p<0.0001), while we observed no significant effect of stimulus type (χ^2^=0.0004, DF=1, p=0.9844) nor of the interaction between the two predictors (χ^2^=0.2768, DF=2, p=0.8707). The post-hoc tests expectedly revealed that spiders responded less to static than to slowly moving stimuli (odds ratio=0.592, SE=0.0765, z=-4.061, p=0.0003), and less to the latter than to the fast-moving ones (odds ratio=0.563, SE=0.0995, z=-3.252, p=0.0069). In contrast, no overall difference was appreciable between the illusory and the control stimuli (odds ratio=0.996, SE=0.0969, z=-0.039, p=1), nor for the static (p=1), slowly moving (p=1), or fast-moving stimuli (p=1).

In the second model, the analysis of deviance revealed a small effect of stimulus type (none, illusory or control; χ^2^=6.1031, DF=2, p=0.04729). In the post-hoc however we observed no difference between no-stimulus and control (odds ratio=1.25, SE=0.141, z=1.996, p=0.0918) nor between no-stimulus and illusion (odds ratio=1.23, SE=0.137, z=1.858, p=0.1264). The effect was probably too small and was removed by the correction.

In the third model we observed no effect of stimulus type, either alone (χ^2^=0.0414, DF=1, p=0.83875) nor in the interaction together with motion speed (χ^2^=0.437, DF=1, p=0.50859), rotation direction (χ^2^=0.2431, DF=1, p=0.62194) nor in the triple interaction (χ^2^=0.0168, DF=1, p=0.89688). Also, similarly to the first model, we observed an effect of motion speed (χ^2^=5.8227, DF=1, p=0.01582). Crucially, we observed an effect of the rotation direction (χ^2^=5.9841, DF=1, p=0.01444) and an effect of its interaction with motion speed (χ^2^=5.0212, DF=1, p=0.02504). Therefore, there was indeed a difference between clockwise and counter-clockwise rotating stimuli, but it seemed to be apparent in control as much as in illusion stimuli. The post-hoc tests revealed that the spiders detected with a higher probability counter-clockwise over clockwise rotating stimuli when both at a slow speed (odds ratio=0.735, SE=0.0684, z=-3.309, p=0.0028), while there was no difference between the two directions in the fast-moving stimuli (odds ratio=0.99, SE=0.0941, z=-0.101, p=1).

### Experiment 2 – Concentric peripheral drift illusion

In experiment 2, we strengthened the illusion by presenting the stimuli as 5 concentric rings, increasing the amount of sudden luminosity changes.

In the first model, we again observed a strong effect of motion speed (GLMM analysis of deviance, χ^2^=156.7548, DF=2, p<0.0001), but no effect of stimulus type (χ^2^=1.0076, DF=1, p=0.3155) nor of the interaction between the two (χ^2^=1.3783, DF=2, p=0.502). As for experiment 1, the post-hoc tests revealed no difference between control and illusory stimuli overall (odds ratio=0.924, SE=0.0849, z=-0.864, p=1), nor for the static (p=1), slowly moving (p=1), or fast-moving stimuli (p=1). Moreover, spiders had a higher chance of responding to slowly moving stimuli than to static ones (odds ratio=0.411, SE=0.0528, z=-6.921, p<0.0001), and a higher chance of responding to fast moving stimuli over slowly moving ones (odds ratio=0.611, SE=0.1061, z=-2.839, p=0.0272).

In the second model, the analysis of deviance revealed a small effect of stimulus type (none, illusory or control; χ^2^=14.143, DF=2, p=0.0008491). In the post-hoc tests we observed no difference between no-stimulus and control (odds ratio=0.918, SE=0.1257, z=-0.628, p=1) but a significant difference between no-stimulus and illusion (odds ratio=0.699, SE=0.0752, z=-3.33, p=0.0017). This was unexpected, as model 1 suggested no difference between illusion and control conditions. Indeed, the response probability between these two conditions were quite similar (control: 21.2%, illusion: 22.7%), in both cases higher than the stimulus-absence reaction probability (17%).

In the third model, we observed a significant effect of the interaction between stimulus type and rotation direction (χ^2^=14.229, DF=1, p=0.0002), and an effect of the triple interaction between stimulus type, motion speed and rotation direction (χ^2^=6.9545, DF=1, p=0.0084). We observed no effect of the other predictors nor interactions. The post-hoc tests revealed that while clockwise and counter-clockwise rotation were not different in the response probability for control stimuli (slow: odds ratio=0.539, SE=0.262, z=2.060, p=0.2757, fast: odds ratio=0.146, SE=0.223, z=0.653, p=1), they were for slowly moving illusory stimuli (odds ratio=-1.114, SE=0.26, z=-4.285, p=0.0001), in the direction of counter-clockwise rotation. There was no difference for fast moving illusory stimuli (odds ratio=-0.213, SE=0.234, z=-0.907, p=1). Note that the illusion is supposedly perceived as rotating clockwise (Agrochao et al., 2020; Faubert & Herbert, 1999; Hsieh et al., 2006), while the most responded actual rotation direction was here counter-clockwise.

### Experiment 3 – Reverse direction illusion

In the third experiment, we tested whether mirroring the illusion stimulus, and therefore supposedly flipping the illusory rotation direction, will also reverse the effect of different rotation directions observed in experiment 2.

The first model showed the same effects as in experiment 2. There was an effect of motion speed (GLMM analysis of deviance, χ^2^=17.723, DF=1, p<0.0001), no effect of stimulus type (χ^2^=1.0004, DF=1, p=0.3172) nor of the interaction between the two (χ^2^=1.5297, DF=1, p=0.2162). The post-hoc tests revealed no difference between static control and illusory stimuli conditions (odds ratio=0.757, SE=0.1342, z=-1.57, p=0.2328). Spiders had a higher chance of responding to slowly moving stimuli with respect to static ones (odds ratio=0.578, SE=0.0769, z=-4.118, p=0.0001).

In the second model, the analysis of deviance revealed no effect of stimulus type (none, illusory or control; χ^2^=5.2094, DF=2, p=0.07393). We still performed the post-hoc tests for completeness. We observed no difference between no-stimulus and control (odds ratio=1.36, SE=0.215, z=1.951, p=0.1022) but a significant difference between no-stimulus and illusion conditions (odds ratio=1.28, SE=0.196, z=1.592, p=0.2228).

In the third model, we observed no effect of stimulus type (χ^2^=2.4979, DF=1, p=0.114), an effect of rotation direction (χ^2^=11.5823, DF=1, p=0.00066), and an effect of the interaction between the two (χ^2^=8.2039, DF=1, p=0.00418). The post-hoc tests revealed that there was no difference between clockwise and counter-clockwise rotating control stimuli (odds ratio=1.09, SE=0.29, z=0.316, p=1). Instead, we observed a higher reaction probability for clockwise over counter-clockwise illusory stimuli (odds ratio=3.14, SE=0.809, z=4.438, p<0.0001), the reverse of experiment 2.

## Discussion

In this paper, we presented a set of tests aimed at discovering whether jumping spiders can perceive the peripheral drift illusion. We presented the animals with stimuli that should induce the perception of illusory motion and stimuli that should not. These stimuli could either appear and remain static, move slowly or move rapidly. In the moving conditions, the objects could rotate either clockwise or counter-clockwise, respectively congruently or in-congruently. We measured the probability that the spider would turn towards the stimulus upon its appearance.

### Jumping spiders do not perceive the peripheral drift illusion

In all 3 experiments, the spiders never reacted to the appearance of static stimuli, being them illusion inducing or control. As expected instead, the spiders responded to both stimuli as soon as these started moving – whether slow or fast. This demonstrates that the spiders in this setup indeed respond to rotating stimuli, but not to static ones. The absence of difference in response probability between the two static stimuli suggests that, as hypothesized, the spiders do not perceive the peripheral drift illusion.

### Peripheral drift stimuli orientation affects detection, but only while rotating

The detection probabilities observed in response to slowly moving stimuli are of more complex interpretation. In experiment 1, spiders showed a higher probability of responding to both the illusory and the control stimuli moving counter-clockwise. In experiment 2 in contrast, this higher probability was observed only for illusory stimuli. The results of experiment 1 especially suggest a general bias for counter-clockwise rotations, which may be more easily detected by the spiders. However, the lack of the same effect in control stimuli of experiment 2 does not fully confirm this explanation. Therefore, in experiment 3 we presented the illusory stimuli flipped, to test whether the “preferred” rotation direction depended on the stimuli itself. As expected, the spiders responded with a higher probability to clockwise rotating illusory stimuli, with no preferences in the control. This demonstrates that the clockwise or counter-clockwise higher detection probability must be driven by the illusory stimulus orientation, rather than an innate tendency of the spiders. We can safely assume that the effect observed in experiment 1 for control stimuli was a priming effect: after detecting the illusory stimuli for a certain number of times, the spiders generalized the same percept to the control ones. This did not occur in experiment 2 and 3, probably due use of more conspicuous stimuli, which made generalizations harder. We never observed any effect of rotation direction in rapidly moving stimuli. In these cases, the high salience of the motion likely masked any other subtle effect generated by the stimuli appearance.

While it is apparent that the illusory stimulus is driving the difference in detection probabilities, the underlying mechanism remains unclear. As stated in the introduction and methods sections, the peripheral drift illusory stimuli we presented in experiment 1 and 2 should induce the perception of a clockwise rotation. Conversely, the stimulus presented in experiment 3 should induce the perception of a counter-clockwise rotation. This observation is valid across taxa. If this were the case for spiders as well, we would expect an increased detection rate for clockwise rotating stimuli in experiment 1 and 2, an increased detection rate for counterclockwise rotating stimuli in experiment 3. In these cases, in fact, the illusory percept should be congruent with motion direction, producing an additive effect in the total perceived speed of rotation. Vice-versa, we should expect a subtraction effect in incongruence combinations (clockwise illusion when the stimuli move counterclockwise), decreasing consequently the detection rate. We observed instead the exact opposite.

### Improbability of a reversed illusion effect in jumping spiders

It may be that jumping spiders do perceive the illusion, but in an opposite direction with respect to all other animals. However, depending on the mechanism that drives the illusion perception, this reverse direction may not be a possibility. As described in the Introduction, the peripheral drift illusion may be generated by a faster processing of higher luminance areas of the visual field compared to darker ones, that when arranged in circular stimuli generates the perception of motion (Conway et al., 2005; Faubert & Herbert, 1999). This explanation depends on the actual luminance differences perceived by various photo-receptors and by physical limitations, making it irreversible (i.e., lower luminance being processed faster). On the other hand, Agrochao et al. (2020) proposed that the illusion percept is caused by an unbalance in the neural circuit itself, with T4 neurons producing a stronger output than T5 neurons in *Drosophila*. If jumping spiders have an analogous division between an ON and OFF edge systems, it could be possible for it to be unbalanced in reverse, with T5 analogous neurons producing a stronger output, thus reversing the effect. However, this explanation is unlikely, as it cannot account for the lack of difference in the detection probability between control and illusory stimuli when presented statically. If the illusory motion generated a strong enough effect to modify the perceived rotation speed, it should have produced an appreciable effect on the static stimuli as well.

### Animacy perception may drive the animals’ response

We propose that the different response probabilities for the different rotation directions are not an effect of additive motion perception (real and illusory). Rather, they are linked to a violation of expectation in the stimuli’ “behavior”. Even if the illusory stimuli in our experiments were not perceived as in motion, they remain asymmetric, and as such their geometry may suggest a natural orientation. It has been shown that many animals assume the natural motion direction of an object in view based solely on its geometry (Rosa-Salva et al., 2018). This shape/motion association is part of the many perceptual rules that allow animals to visually detect other living organisms and distinguish them from inanimate in their visual field (A skill termed “animacy perception”. Brown et al., 2010; Johnson, 2006; Rosa-Salva et al., 2016; Vallortigara, 2021). For example, all *bilateria* tend to move in a direction parallel, rather than perpendicular, to their major body axis (Rosa-Salva et al., 2018). We already know that jumping spiders are equipped with at least some of these perceptual rules (De Agrò et al., 2021), and that generally respond more to stimuli that do not follow them. We hypothesize that the peripheral drift illusion stimuli, although not perceived as in motion, do suggest a natural predisposition to clockwise or counter-clockwise motion direction, leading to a higher response rate when not respected.

### What can the lack of illusory percept tell us about jumping spiders

The absence of illusory perception in the peripheral drift illusion in jumping spiders can provide insight into how their visual system functions. On one hand, it is possible that motion perception circuitry in the spider’s secondary eyes is structured similarly to what has been observed in *Drosophila* and other insects: divided into two streams for light and dark edges, parallel computing the direction of motion across neighboring photoreceptors (Reichardt, 1987). If this is the case however, their system must lack the imbalance between the ON and OFF circuits observed by Agrochao et. al. (2020). On the other hand, spiders may perceive motion without relying on the parallel circuits for light and dark edges. As stated in the introduction, the unique motion detection system of these animals is divided into 3 functionally specialized eye-pairs. It is possible that the ON and OFF edge systems are also distributed across different eyes, which would render any imbalance between the two circuits irrelevant, since given their distinct visual fields they would never perceive the same stimulus simultaneously.

Regardless of which interpretation we lean towards, the results we observed remain surprising. Illusory motion perception seems to be ubiquitous across animals, along with its underlying neural circuitry. Moreover, it has been observed that the genes regulatory network governing the development of compound eyes in *Drosophila* have orthologs in at least two spider species (*Parasteatoda tepidariorum* and *Cupiennius salei*) (N. Morehouse, 2020; N. I. Morehouse et al., 2017), demonstrating a deep conservation of the eyes across taxa. We propose then that the different perceptual responses between *Drosophila* and jumping spiders must have arisen because the visual system of spiders is split in multiple eyes. Future studies could test whether ON and OFF edge system are indeed divided across separate eye-pairs, and what effects this split generates.

In this study, we tested the perception of motion in the peripheral drift illusion by a jumping spider. We observed that while detecting the stimulus orientation, the spiders did not perceive it as moving. We discussed the implications of this findings both for the mechanistic hypothesis explaining the illusory percept and for the spider’s visual system itself. Our observation contributes to the growing literature on the specialization of secondary eyes in jumping spiders, highlighting the uniqueness of their visual system across the animal kingdom.

## Methods

### Subjects

We employed a total of 117 *Menemerus semilimbatus* (77 females, 18 males and 22 juveniles). All animals were collected in the wild, in Florence, Italy, between April and May 2023. Once caught, we applied to the cephalothorax a cylindrical, 1×1mm neodymium magnet, using UV activated resin. This magnet was needed to fix the animal to the apparatus (see the next section). The spiders were then housed individually in plastic boxes and tested for the first time the next day. At the end of the experiment, the magnet was removed and the subject was released back into nature.

### Overall procedure

All the experiments were performed using a spherical treadmill as described in (De Agrò et al., 2021). Briefly, each spider was attached to the end effector of a 6-axis micro-manipulator, thanks to the neodymium magnet. Then, the animal was placed above and in contact with a polystyrene sphere (39mm in diameter). This ball was made friction-less thanks to a constant stream of compressed air from below. In this setup, the spider was unable to shift its position due to the magnet, but was still able to move its legs and impress its intended motion onto the sphere. By measuring the sphere rotation, we were able to infer the animal intended turning direction.

The full setup was placed in front of a computer monitor, at 112mm distance from it. The monitor displayed a white noise pattern as background. After 210 seconds of acclimatization to become accustomed with the setup, the experiment would start.

Either to the left or to the right of the spider, at a distance of 40° from the center, a stimulus appeared. As previously stated, jumping spiders produce typical whole-body pivots upon detection of a moving stimulus by the secondary eyes, in order to put it in view of the principal eyes. Being presented at ±40°, we can be sure that the stimuli were presented in the ALEs visual fields, but outside of the AMEs ones (De Agrò et al., 2024). When the spider attempted a pivot towards the stimulus, we observed a counter-rotation of the polystyrene sphere, allowing us to infer the animals’ perception for different stimuli. Each stimulus remained on screen for 2 seconds, then disappeared. After 20 seconds, the next stimulus was presented, for a total of 30 stimuli. Each animal underwent a single trial.

### Experiment 1 – Annular peripheral drift illusion

In the first experiment, we tested the ability of the spiders to detect the peripheral drift illusion as a moving object. In order to do so we designed 2 different stimuli (Figure 1), and each one could move in one of three modalities. The first stimulus (annular illusion) displayed a saw-tooth gradient with 20 cycles per rotation, going from light to dark in the clockwise direction. The stimulus was circular in shape and 20° wide. The saw-tooth pattern was however only displayed for 1/3rd of the full circle, leaving the center empty, creating an annular pattern. Given the orientation of the gradient, this stimulus should induce the perception of a clockwise rotation (Agrochao et al., 2020; Faubert & Herbert, 1999; Hsieh et al., 2006).

The second stimulus acted as a control (annular control). This was identical to the annular illusion, but presented a sine-wave pattern rather than a saw-tooth one: going from dark to light and from light to dark, the stimulus appears symmetrical, with no presence of hard changes. This stimulus while maintaining all low-level characteristics of the previous one (contrast, luminance, etc.), should not induce any illusory motion.

Upon appearance on the monitor, the stimuli could produce one of three behaviors:

1. appear to the left or right of the spider, remain static for the full 2 seconds of visibility, then disappear
2. appear to the left or right of the spider, rotate clockwise or counter-clockwise around its center at a speed of 2° per second, disappear.
3. appear to the left or right of the spider, rotate clockwise or counter-clockwise around its center at a speed of 32° per second, disappear.

If the spiders perceive the peripheral drift illusion as moving, we hypothesize to observe no reaction to the appearing annular control stimulus (we expect stimulus appearance to not be enough to trigger a pivot, consistent with observations in Ferrante et al., 2023), but a response to the annular illusion stimulus, possibly equal to the slowly moving control. For the slowly moving stimuli, we expect to observe instead an equal reaction for annular controls, independently of rotation direction, but a difference for the annular illusion, depending on whether the stimulus motion was congruent or in-congruent to the illusion direction. The fast-moving stimuli are included as a secondary control, to check whether the stimulus speed has an effect on pivot probability.

A total of 46 spiders underwent experiment 1. Each animal was subjected to all stimuli: position on the monitor, stimulus type, stimulus speed and motion direction were all determined randomly for each of the 30 stimuli in the trial.

### Experiment 2 – Concentric peripheral drift illusion

Having observed no effect of the illusory stimuli in experiment 1 (see results), we performed a second experiment. It is possible that the presence of only 1 gradient ring in the illusion, although present, was insufficient to induce a strong enough effect to elicit a pivot in the spider. For this reason, we designed two new stimuli (Concentric illusion and concentric control, Figure 1). Here, the saw-tooth and sine-wave gradients were repeated as concentric rings, rotated 9° with respect to each other. This caused the presence of 5 times as many hard changes from light to dark in the illusory stimulus, supposedly multiplying the strength of the illusory effect. For the control condition, in contrast, no difference should be appreciable, if not an increased number of pixel changes upon appearance and motion of the object. The stimuli were presented as in the experiment 1, being either static, slow or fast moving, clockwise or counter-clockwise. A total of 48 spiders underwent experiment 2. Each animal was subjected to all stimuli: position on the monitor, stimulus type, stimulus speed and motion direction were all determined randomly for each of the 30 stimuli in the trial.

### Experiment 3 – Reverse direction illusion

Having observed a counter-intuitive effect of rotation direction for the illusory stimuli upon detection probability (see results and discussion), we repeated experiment 2 after flipping the illusory stimuli (Figure 1). With this adjustment, if related to the saw-tooth gradient orientation, we expected that any motion direction effect should have reversed. For this experiment, we only included the slowly moving stimuli (2° per second), being the rotation direction effect prevalent only at this speed in experiment 2. A total of 23 spiders underwent experiment 2. Each animal was subjected to all stimuli: position on the monitor, stimulus type, stimulus speed and motion direction were all determined randomly for each of the 30 stimuli in the trial.

### Scoring

The rotations of the spherical treadmill, collected with the software FicTrac (Moore et al., 2014), were further analysed through an algorithm developed in Python 3.10 (Van Rossum & Drake, 2009), available as a supplement (SI2). This algorithm extracts every rotation of the sphere around its Z axis of at least a total of 10° with no concurrent rotations around the X or Y axes. Moreover, these rotations had to last between 0.1 and 1.5 seconds in total. This movement pattern corresponds to the spider pivoting towards the moving stimulus detected by secondary eyes. For every stimulus presented, we observed whether for its whole duration any pivot was produced, and if it was congruent with the stimulus direction. If all the aforementioned requirements were met, we considered the stimulus to have been detected, and a pivot produced.

We also performed the same procedure for random time-points during the inter-stimulus interval, so when no change on stimulus happened. This was done to collect a baseline value of the animals’ reaction in the complete absence of moving targets. Indeed, while the observed rotations in this case should ideally be 0, the presence of noise in the system and spontaneous activity of the animal will cause the registering of some responses. This value will be useful for determining whether the appearance of static stimuli can cause any response.

### Statistical analysis

All analyses were performed in R4.1.2 (R Core Team, 2021). Raw data is available in SI1. Analysis script is available in SI2. For each experiment, we performed 3 separate generalized mixed effect models using the package glmmTMB (Bolker et al., 2009; Brooks et al., 2017), all with binomial error structure, to test the effect of various predictors on detection probability. In the first model, we tested the effect of stimulus type (illusory or control), of motion speed (static, slow or fast) and the interaction between the two. In the second model, we only included static stimuli and tested them against no-stimulus presentation. In the third, we only included moving stimuli, to test the effect of rotation direction. In all three models we included subject identity as a random effect. The goodness of fit was tested using the package DHARMa (Hartig, 2018). We then performed an analysis of deviance using the package car (Fox & Weisberg, 2019), to test which of the predictors had an effect on the dependent variable. We then performed Bonferroni corrected post-hoc tests using the package emmeans (Lenth et al., 2020). Lastly, using the package reticulate (Ushey et al., 2021) to communicate with python 3.10 (Van Rossum & Drake, 2009), we produced plots with the libraries pandas (Reback et al., 2020) matplotlib (Hunter, 2007) and seaborn (Waskom et al., 2017).

## Acknowledgements

We would like to thank Livia De Fazi for the help with animal collection. MDA was partially funded by a grant by “Fondazione Cassa Di Risparmio Di Trento e Rovereto”

## References

Agrillo, C., Gori, S., & Beran, M. J. (2015). Do rhesus monkeys (Macaca mulatta) perceive illusory motion? Animal Cognition, 18(4), 895–910. 10.1007/s10071-015-0860-6

Agrochao, M., Tanaka, R., Salazar-Gatzimas, E., & Clark, D. A. (2020). Mechanism for analogous illusory motion perception in flies and humans. Proceedings of the National Academy of Sciences, 117(37), 23044–23053. 10.1073/pnas.2002937117

Bååth, R., Seno, T., & Kitaoka, A. (2014). Cats and Illusory Motion. Psychology, 2014. 10.4236/psych.2014.59125

Beydizada, N. I., Cannone, F., Pekár, S., Baracchi, D., & De Agrò, M. (2024). Habituation to visual stimuli is independent of boldness in a jumping spider. Animal Behaviour, 213, 61–70. 10.1016/j.anbehav.2024.04.010

Bolker, B. M., Brooks, M. E., Clark, C. J., Geange, S. W., Poulsen, J. R., Stevens, M. H. H., & White, J.-S. S. (2009). Generalized linear mixed models: A practical guide for ecology and evolution. Trends in Ecology & Evolution, 24(3), Article 3. 10.1016/j.tree.2008.10.008

Brooks, M. E., Kristensen, K., Benthem, K. J. van, Magnusson, A., Berg, C. W., Nielsen, A., Skaug, H. J., Maechler, M., & Bolker, B. M. (2017). glmmTMB Balances Speed and Flexibility Among Packages for Zero-inflated Generalized Linear Mixed Modeling. The R Journal, 9(2), 378–400.

Brown, J., Kaplan, G., Rogers, L. J., & Vallortigara, G. (2010). Perception of biological motion in common marmosets (Callithrix jacchus): By females only. Animal Cognition, 13(3), 555–564. 10.1007/s10071-009-0306-0

Bruce, M., Daye, D., Long, S. M., Winsor, A. M., Menda, G., Hoy, R. R., & Jakob, E. M. (2021). Attention and distraction in the modular visual system of a jumping spider. Journal of Experimental Biology, 224(8). 10.1242/jeb.231035

Chong, K. L., Grahn, A., Perl, C. D., & Sumner-Rooney, L. (2024). Allometry and ecology shape eye size evolution in spiders. Current Biology, 0(0). 10.1016/j.cub.2024.06.020

Conway, B. R., Kitaoka, A., Yazdanbakhsh, A., Pack, C. C., & Livingstone, M. S. (2005). Neural Basis for a Powerful Static Motion Illusion. Journal of Neuroscience, 25(23), 5651–5656. 10.1523/JNEUROSCI.1084-05.2005

De Agrò, M., Rößler, D. C., Kim, K., & Shamble, P. S. (2021). Perception of biological motion by jumping spiders. PLOS Biology, 19(7), Article 7. 10.1371/journal.pbio.3001172

De Agrò, M., Rößler, D. C., & Shamble, P. S. (2024). Eye-specific detection and a multi-eye integration model of biological motion perception. Journal of Experimental Biology, jeb.247061. 10.1242/jeb.247061

Dolev, Y., & Nelson, X. J. (2014). Innate pattern recognition and categorization in a jumping spider. PloS One, 9(6), e97819. 10.1371/journal.pone.0097819

Eagleman, D. M. (2001). Visual illusions and neurobiology. Nature Reviews Neuroscience, 2(12), Article 12. 10.1038/35104092

Faubert, J., & Herbert, A. M. (1999). The Peripheral Drift Illusion: A Motion Illusion in the Visual Periphery. Perception, 28(5), 617–621. 10.1068/p2825

Ferrante, F., Loconsole, M., Giacomazzi, D., & Agrò, M. D. (2023). Separate attentional processes in the two visual systems of jumping spiders (p. 2023.04.13.536553). bioRxiv. 10.1101/2023.04.13.536553

Fox, J., & Weisberg, S. (2019). An R Companion to Applied Regression (Third). Sage. https://socialsciences.mcmaster.ca/jfox/Books/Companion/

Fraser, A., & Wilcox, K. J. (1979). Perception of illusory movement. Nature, 281(5732), Article 5732. 10.1038/281565a0

Gori, S., Agrillo, C., Dadda, M., & Bisazza, A. (2014). Do Fish Perceive Illusory Motion? Scientific Reports, 4(1), Article 1. 10.1038/srep06443

Hartig, F. (2018). DHARMa: Residual Diagnostics for Hierarchical (Multi-Level / Mixed) Regression Models. https://CRAN.R-project.org/package=DHARMa

Hsieh, P.-J., Caplovitz, G. P., & Tse, P. U. (2006). Illusory motion induced by the offset of stationary luminance-defined gradients. Vision Research, 46(6), 970–978. 10.1016/j.visres.2005.10.009

Hunter, J. D. (2007). Matplotlib: A 2D graphics environment. Computing in Science & Engineering, 9(3), 90–95.

Jakob, E. M., Long, S. M., Harland, D. P., Jackson, R. R., Carey, A., Searles, M. E., Porter, A. H., Canavesi, C., & Rolland, J. P. (2018). Lateral eyes direct principal eyes as jumping spiders track objects. Current Biology, 28(18), R1092–R1093. 10.1016/j.cub.2018.07.065

Johnson, M. H. (2006). Biological Motion: A Perceptual Life Detector? Current Biology, 16(10), R376–R377. 10.1016/j.cub.2006.04.008

Kanizsa, G. (1979). Organization in vision: Essays on gestalt perception. Praeger.

Kelley, L. A., & Kelley, J. L. (2014). Animal visual illusion and confusion: The importance of a perceptual perspective. Behavioral Ecology, 25(3), 450–463. 10.1093/beheco/art118

Kitaoka, A. (2003). Rotating snakes. http://www.Ritsumei.Ac.Jp/~Akitaoka/Index-e.Html.

Kitaoka, A., & Ashida, H. (2003). Phenomenal Characteristics of the Peripheral Drift Illusion. Vision, 15(4), 261–262. 10.24636/vision.15.4_261

Land, M. F. (1969). Movements of the retinae of jumping spiders (Salticidae: Dendryphantinae) in response to visual stimuli. The Journal of Experimental Biology, 51(2), 471–493.

Land, M. F. (1985). Short Communication: Fields of View of the Eyes of Primitive Jumping Spiders. Journal of Experimental Biology, 119(1), 381–384.

Lazareva, O. F., Shimizu, T., & Wasserman, E. A. (2012). How Animals See the World: Comparative Behavior, Biology, and Evolution of Vision. Oxford University Press. 10.1093/acprof:oso/9780195334654.001.0001

Lenth, R., Singmann, H., Love, J., Buerkner, P., & Herve, M. (2020). emmeans: Estimated Marginal Means, aka Least-Squares Means (Version 1.4.5) [Computer software]. https://CRAN.R-project.org/package=emmeans

Moore, R. J. D., Taylor, G. J., Paulk, A. C., Pearson, T., van Swinderen, B., & Srinivasan, M. V. (2014). FicTrac: A visual method for tracking spherical motion and generating fictive animal paths. Journal of Neuroscience Methods, 225, 106–119. 10.1016/j.jneumeth.2014.01.010

Morehouse, N. (2020). Spider vision. Current Biology, 30(17), R975–R980.

Morehouse, N. I., Buschbeck, E. K., Zurek, D. B., Steck, M., & Porter, M. L. (2017). Molecular Evolution of Spider Vision: New Opportunities, Familiar Players. The Biological Bulletin, 233(1), 21–38. 10.1086/693977

Murakami, I., Kitaoka, A., & Ashida, H. (2006). A positive correlation between fixation instability and the strength of illusory motion in a static display. Vision Research, 46(15), 2421–2431. 10.1016/j.visres.2006.01.030

Nieder, A. (2002). Seeing more than meets the eye: Processing of illusory contours in animals. Journal of Comparative Physiology A, 188(4), 249–260. 10.1007/s00359-002-0306-x

Otero-Millan, J., Macknik, S. L., & Martinez-Conde, S. (2012). Microsaccades and Blinks Trigger Illusory Rotation in the “Rotating Snakes” Illusion. Journal of Neuroscience, 32(17), 6043–6051. 10.1523/JNEUROSCI.5823-11.2012

R Core Team. (2021). R: A Language and Environment for Statistical Computing. R Foundation for Statistical Computing. https://www.R-project.org/

Reback, J., McKinney, W., jbrockmendel, Bossche, J. V. den, Augspurger, T., Cloud, P., gfyoung, Sinhrks, Klein, A., Roeschke, M., Hawkins, S., Tratner, J., She, C., Ayd, W., Petersen, T., Garcia, M., Schendel, J., Hayden, A., MomIsBestFriend, … Mehyar, M. (2020). pandas-dev/pandas: Pandas 1.0.3 [Computer software]. Zenodo. 10.5281/zenodo.3715232

Reichardt, W. (1987). Evaluation of optical motion information by movement detectors. Journal of Comparative Physiology A, 161(4), 533–547. 10.1007/BF00603660

Rosa Salva, O., Sovrano, V. A., & Vallortigara, G. (2014). What can fish brains tell us about visual perception? Frontiers in Neural Circuits, 8. 10.3389/fncir.2014.00119

Rosa-Salva, O., Grassi, M., Lorenzi, E., Regolin, L., & Vallortigara, G. (2016). Spontaneous preference for visual cues of animacy in naïve domestic chicks: The case of speed changes. Cognition, 157, 49–60. 10.1016/j.cognition.2016.08.014

Rosa-Salva, O., Hernik, M., Broseghini, A., & Vallortigara, G. (2018). Visually-naïve chicks prefer agents that move as if constrained by a bilateral body-plan. Cognition, 173, 106–114. 10.1016/j.cognition.2018.01.004

Rößler, D. C., De Agrò, M., Kim, K., & Shamble, P. S. (2022). Static visual predator recognition in jumping spiders. Functional Ecology, 36(3), 561–571. 10.1111/1365-2435.13953

Strausfeld, N. J., & Barth, F. G. (1993). Two visual systems in one brain: Neuropils serving the secondary eyes of the spider Cupiennius salei. Journal of Comparative Neurology, 328(1), 43–62. 10.1002/cne.903280104

Strausfeld, N. J., Weltzien, P., & Barth, F. G. (1993). Two visual systems in one brain: Neuropils serving the principal eyes of the spider Cupiennius salei. Journal of Comparative Neurology, 328(1), 63–75. 10.1002/cne.903280105

Todorović, D. (2020). What Are Visual Illusions?*. Perception, 49(11), 1128–1199. 10.1177/0301006620962279

Ushey, K., Allaire, J. J., & Tang, Y. (2021). reticulate: Interface to ‘Python’. https://CRAN.R-project.org/package=reticulate

Vallortigara, G. (2004). Visual Cognition and Representation in Birds and Primates. In L. J. Rogers & G. Kaplan (Eds.), Comparative Vertebrate Cognition: Are Primates Superior to Non-Primates? (pp. 57–94). Springer US. 10.1007/978-1-4419-8913-0_2

Vallortigara, G. (with Losi, C.). (2021). Born Knowing: Imprinting and the Origins of Knowledge. The MIT Press.

Van Rossum, G., & Drake, F. L. (2009). Python 3 Reference Manual. CreateSpace.

Waskom, M., Botvinnik, O., O’Kane, D., Hobson, P., Lukauskas, S., Gemperline, D. C., Augspurger, T., Halchenko, Y., Cole, J. B., Warmenhoven, J., Ruiter, J. de, Pye, C., Hoyer, S., Vanderplas, J., Villalba, S., Kunter, G., Quintero, E., Bachant, P., Martin, M., … Qalieh, A. (2017). mwaskom/seaborn: V0.8.1 (September 2017) (Version v0.8.1) [Computer software]. Zenodo. 10.5281/zenodo.883859

Winsor, A. M., Pagoti, G. F., Daye, D. J., Cheries, E. W., Cave, K. R., & Jakob, E. M. (2021). What gaze direction can tell us about cognitive processes in invertebrates. Biochemical and Biophysical Research Communications, 564, 43–54. 10.1016/j.bbrc.2020.12.001

Winsor, A. M., Remage-Healey, L., Hoy, R. R., & Jakob, E. M. (2024). Visual attention and processing in jumping spiders. Trends in Neurosciences, 47(1), 6–8. 10.1016/j.tins.2023.09.002

Zurek, D. B., & Nelson, X. J. (2012a). Hyperacute motion detection by the lateral eyes of jumping spiders. Vision Research, 66, 26–30. 10.1016/j.visres.2012.06.011

Zurek, D. B., & Nelson, X. J. (2012b). Saccadic tracking of targets mediated by the anterior-lateral eyes of jumping spiders. Journal of Comparative Physiology A, 198(6), 411–417. 10.1007/s00359-012-0719-0

Zurek, D. B., Taylor, A. J., Evans, C. S., & Nelson, X. J. (2010). The role of the anterior lateral eyes in the vision-based behaviour of jumping spiders. Journal of Experimental Biology, 213(14), 2372–2378. 10.1242/jeb.042382

