## Supplementary material for "Jumping spiders are not fooled by the peripheral drift illusion": SI2.html


Code 

- Show All Code
- Hide All Code

### SI2 - Analysis | Jumping spiders are not fooled by the peripheral drift illusion

#### SI2 - Analysis | Jumping spiders are not fooled by the peripheral drift illusion

- 1 Prepare environment
  - 1.1 loading packages
  - 1.2 loading files
  - 1.3 Prepare tables
- 2 Analysis
  - 2.1 Preliminary questions
    - 2.1.1 Describe subjects
    - 2.1.2 Is there an habituation or fatigue effect?
    - 2.1.3 Differences between sexes
  - 2.2 Main analysis
    - 2.2.1 Experiment 1 - ring stimuli
    - 2.2.2 Experiment 2 - concentric stimuli
    - 2.2.3 Experiment 3 - flipped concentric stimuli

2024-09-17

Massimo De Agrò (1,2,3), Giorgio Vallortigara (3), Egidio Falotico (1,2)

1 – The BioRobotics Institute, Sant’Anna School of Advanced Studies, Pontedera (PI), Italy

2 – Department of Excellence in Robotics, Sant’Anna School of Advanced Studies, Pontedera (PI), Italy

3 – Center for Mind-Brain Sciences, University of Trento, Rovereto, Italy

---

This document (SI2) contains the statistical analysis output for the manuscript. SI1 provides the entire raw data on which this analysis is based. Note that much of the analysis is divided into sections using tabs, found at the start of the sections. Click on the section you wish to inspect, and scroll down for the full analysis.

---

In this experiment, we tested the ability of jumping spiders to perceive the peripheral drift illusion. We hypothesize that these animals will not be fooled by the illusion, as they supposedly lack the required neural substrate.

We designed 4 different experimental conditions:

- peripheral drift, illusion inducing stimulus, static
- peripheral drift, illusion inducing stimulus, rotating
  - either clockwise
  - or counterclockwise
  - either slowly (2 deg per second)
  - or fast (32 deg per second)
- control stimulus, not illusion inducing, static
- control stimulus, not illusion inducing, rotating
  - either clockwise
  - or counterclockwise
  - either slowly (2 deg per second)
  - or fast (32 deg per second)

We performed 2 experiments. In the first, the control and illusory stimuli were presented as a “ring”. In the second, 5 concentric rings were presented. This was done to enhance the illusory effect, given the increased number of color-change areas.

### 1 Prepare environment

#### 1.1 loading packages

```
library(data.table) #To deal with big datasets
library(glmmTMB)
library(emmeans)
library(car)
```

```
## Loading required package: carData
```

```
library(DHARMa)
```

```
## This is DHARMa 0.4.6. For overview type '?DHARMa'. For recent changes, type news(package = 'DHARMa')
```

```
library(ggplot2)
library(reticulate)
set.seed(12345)
```

#### 1.2 loading files

```
data <- fread(paste0(path, 'SI1.csv'))
data$date <- as.factor(data$date)
data$subj <- as.factor(data$subj)
data$sex <- as.factor(data$sex)
data$stimn <- as.numeric(data$stimn)
data$side <- as.factor(data$side)
data$angle <- as.numeric(data$angle)
data$stim_type <- as.factor(data$stim_type)
data$speed <- as.factor(data$speed)

data$stim_motion <- as.factor(data$stim_motion)
data$rot_direction <- as.factor(data$rot_direction)
```

#### 1.3 Prepare tables

```
#restructure motion as its own variable, no motion, speed 2 and speed 32
data$motion <- 0
data$motion[data$speed == 2] <- 2
data$motion[data$speed == 32] <- 32

data$motion <- as.factor(data$motion)

#splitting the 3 experimentss
exp1 <- subset(data, data$exp == 'Ring')
exp2 <- subset(data, data$exp == 'Concentric')
exp3 <- subset(data, data$exp == 'Flipped')

# These are for comparisons
exp1_onlystim <-subset(exp1, exp1$stim_type != 'none')
exp2_onlystim <-subset(exp2, exp2$stim_type != 'none')
exp3_onlystim <-subset(exp3, exp3$stim_type != 'none')

# These are for motion analysis
exp1_onlymoving <-subset(exp1_onlystim, exp1_onlystim$stim_motion == 'Moving')
exp2_onlymoving <-subset(exp2_onlystim, exp2_onlystim$stim_motion == 'Moving')
exp3_onlymoving <-subset(exp3_onlystim, exp3_onlystim$stim_motion == 'Moving')

# These are for comparison with nostim
exp1_onlystatic <-subset(exp1, exp1$motion == 0)
exp2_onlystatic <-subset(exp2, exp2$motion == 0)
exp3_onlystatic <-subset(exp3, exp3$motion == 0)
```

### 2 Analysis

#### 2.1 Preliminary questions

##### 2.1.1 Describe subjects

```
# First, let's count
onerow_per_subj <- subset(data, data$stimn==0)
onerow_per_subj <- subset(onerow_per_subj, onerow_per_subj$trialn==1)
summary(as.factor(onerow_per_subj$exp))
```

```
## Concentric    Flipped       Ring 
##         48         23         46
```

```
summary(onerow_per_subj$sex)
```

```
##  f  j  m  M 
## 77 22 18  0
```

unfortunately experiment 3 has fewer subjects. that’s life

##### 2.1.2 Is there an habituation or fatigue effect?

We first want to see if response rate (independently from stimulus type) decreases across trials and stimuli

```
pm_f <- glmmTMB(turn~trialn*stimn + (1|subj), data = data, family = binomial())
simres <- simulateResiduals(pm_f)
plot(simres)
```

```
Anova(pm_f)
```

```
## Analysis of Deviance Table (Type II Wald chisquare tests)
## 
## Response: turn
##                 Chisq Df Pr(>Chisq)    
## trialn         2.6698  1     0.1023    
## stimn        190.5264  1     <2e-16 ***
## trialn:stimn   0.4898  1     0.4840    
## ---
## Signif. codes:  0 '***' 0.001 '**' 0.01 '*' 0.05 '.' 0.1 ' ' 1
```

```
e <- emmeans(pm_f, ~trialn, type='response')
```

```
## NOTE: Results may be misleading due to involvement in interactions
```

```
e #just to see
```

```
##  trialn  prob     SE  df asymp.LCL asymp.UCL
##       1 0.286 0.0174 Inf     0.253     0.322
##       2 0.265 0.0173 Inf     0.233     0.301
## 
## Confidence level used: 0.95 
## Intervals are back-transformed from the logit scale
```

```
et <- emtrends(pm_f, ~1, var = 'stimn', type='response')

test(et)
```

```
##  1       stimn.trend      SE  df z.ratio p.value
##  overall     -0.0474 0.00344 Inf -13.795  <.0001
## 
## Results are averaged over the levels of: trialn
```

No difference between the two trials. However, the probability of responses decreases of around 5% per stimulus

```
ggplot(data, aes(x=stimn, y=turn))+
        geom_smooth(method = 'glm')+
        ylim(0,1)
```

```
## `geom_smooth()` using formula = 'y ~ x'
```

```
## Warning: Removed 6415 rows containing non-finite values (`stat_smooth()`).
```

##### 2.1.3 Differences between sexes

what about the response rate differences between males, females and juveniles?

```
# First, let's count
onerow_per_subj <- subset(data, data$stimn==0)
onerow_per_subj <- subset(onerow_per_subj, onerow_per_subj$trialn==1)

summary(onerow_per_subj$sex)
```

```
##  f  j  m  M 
## 77 22 18  0
```

we mostly have females, which is dependent from availability in nature. The results should be interpreted with caution given the large disparity, but will show.

```
pm_s <- glmmTMB(turn~sex + (1|subj), data = data, family = binomial())
simres <- simulateResiduals(pm_s)
plot(simres)
```

```
Anova(pm_s)
```

```
## Analysis of Deviance Table (Type II Wald chisquare tests)
## 
## Response: turn
##      Chisq Df Pr(>Chisq)
## sex 4.9763  3     0.1735
```

```
e <- emmeans(pm_s, ~sex, type='response')
e #just to see
```

```
##  sex  prob     SE  df asymp.LCL asymp.UCL
##  f   0.233 0.0121 Inf     0.210     0.257
##  j   0.250 0.0225 Inf     0.208     0.296
##  m   0.189 0.0217 Inf     0.151     0.236
##  M   0.255 0.0724 Inf     0.140     0.420
## 
## Confidence level used: 0.95 
## Intervals are back-transformed from the logit scale
```

```
pairs(e)
```

```
##  contrast odds.ratio    SE  df null z.ratio p.value
##  f / j         0.912 0.123 Inf    1  -0.681  0.9045
##  f / m         1.299 0.203 Inf    1   1.675  0.3371
##  f / M         0.885 0.342 Inf    1  -0.316  0.9890
##  j / m         1.424 0.254 Inf    1   1.988  0.1925
##  j / M         0.970 0.384 Inf    1  -0.076  0.9998
##  m / M         0.681 0.249 Inf    1  -1.052  0.7186
## 
## P value adjustment: tukey method for comparing a family of 4 estimates 
## Tests are performed on the log odds ratio scale
```

no differences between the sexes

#### 2.2 Main analysis

Now to the main analysis. How do spiders react to the different stimuli?

##### 2.2.1 Experiment 1 - ring stimuli

###### 2.2.1.1 Difference between stimuli

```
mm_s1 <- glmmTMB(turn~stim_type*motion + (1|subj), data = exp1_onlystim, family = binomial())
simres <- simulateResiduals(mm_s1)
plot(simres)
```

```
Anova(mm_s1)
```

```
## Analysis of Deviance Table (Type II Wald chisquare tests)
## 
## Response: turn
##                    Chisq Df Pr(>Chisq)    
## stim_type         0.0004  1     0.9844    
## motion           80.1041  2     <2e-16 ***
## stim_type:motion  0.2768  2     0.8707    
## ---
## Signif. codes:  0 '***' 0.001 '**' 0.01 '*' 0.05 '.' 0.1 ' ' 1
```

```
e <- emmeans(mm_s1, ~stim_type*motion, type='response')

e
```

```
##  stim_type       motion  prob     SE  df asymp.LCL asymp.UCL
##  Control         0      0.193 0.0242 Inf     0.150     0.245
##  PeripheralDrift 0      0.187 0.0235 Inf     0.145     0.237
##  Control         2      0.276 0.0359 Inf     0.212     0.352
##  PeripheralDrift 2      0.291 0.0370 Inf     0.224     0.369
##  Control         32     0.416 0.0444 Inf     0.332     0.504
##  PeripheralDrift 32     0.411 0.0454 Inf     0.325     0.502
## 
## Confidence level used: 0.95 
## Intervals are back-transformed from the logit scale
```

```
write.csv(e, paste0(imagepath, 'exp1.csv'), row.names = FALSE, dec='.')
```

```
## Warning in write.csv(e, paste0(imagepath, "exp1.csv"), row.names = FALSE, :
## attempt to set 'dec' ignored
```

```
contrast(e, list('speed0vs2'=c(0.5, 0.5, -0.5, -0.5, 0, 0),
                 'speed2vs32'=c(0, 0, 0.5, 0.5, -0.5, -0.5),
                 'controlVsDrift'=c(1/3, -1/3, 1/3, -1/3, 1/3, -1/3),
                 'StaticControlVsDrift'=c(1, -1, 0, 0, 0, 0),
                 'SlowControlVsDrift'=c(0, 0, 1, -1, 0, 0),
                 'FastControlVsDrift'=c(0, 0, 0, 0, 1, -1)),
         adjust = 'bonferroni')
```

```
##  contrast             odds.ratio     SE  df null z.ratio p.value
##  speed0vs2                 0.592 0.0765 Inf    1  -4.061  0.0003
##  speed2vs32                0.563 0.0995 Inf    1  -3.252  0.0069
##  controlVsDrift            0.996 0.0969 Inf    1  -0.039  1.0000
##  StaticControlVsDrift      1.043 0.1506 Inf    1   0.294  1.0000
##  SlowControlVsDrift        0.929 0.1622 Inf    1  -0.423  1.0000
##  FastControlVsDrift        1.020 0.1875 Inf    1   0.110  1.0000
## 
## P value adjustment: bonferroni method for 6 tests 
## Tests are performed on the log odds ratio scale
```

No difference between stimuli type! Nor in motion, nor static. this is the main take home message. Interestingly, but fairly obviously, response probability increase with motion speed.

###### 2.2.1.2 Difference between static and nothing

this can be tested by checking whether the probability of pivot in absence of any visual stimulus is different from when stimuli appear.

```
mm_s1_s <- glmmTMB(turn~stim_type + (1|subj), data = exp1_onlystatic, family = binomial())
simres <- simulateResiduals(mm_s1_s)
plot(simres)
```

```
Anova(mm_s1_s)
```

```
## Analysis of Deviance Table (Type II Wald chisquare tests)
## 
## Response: turn
##            Chisq Df Pr(>Chisq)  
## stim_type 6.1031  2    0.04729 *
## ---
## Signif. codes:  0 '***' 0.001 '**' 0.01 '*' 0.05 '.' 0.1 ' ' 1
```

```
e <- emmeans(mm_s1_s, ~stim_type, type='response')

e
```

```
##  stim_type        prob     SE  df asymp.LCL asymp.UCL
##  Control         0.207 0.0199 Inf     0.170     0.248
##  none            0.172 0.0123 Inf     0.149     0.198
##  PeripheralDrift 0.204 0.0195 Inf     0.168     0.244
## 
## Confidence level used: 0.95 
## Intervals are back-transformed from the logit scale
```

```
write.csv(e, paste0(imagepath, 'exp1_static.csv'), row.names = FALSE, dec='.')
```

```
## Warning in write.csv(e, paste0(imagepath, "exp1_static.csv"), row.names =
## FALSE, : attempt to set 'dec' ignored
```

```
contrast(e, list('ControlVsNoStim'=c(1, -1, 0),
                 'IllusionVsNoStim'=c(0, -1, 1)),
         adjust = 'bonferroni')
```

```
##  contrast         odds.ratio    SE  df null z.ratio p.value
##  ControlVsNoStim        1.25 0.141 Inf    1   1.996  0.0918
##  IllusionVsNoStim       1.23 0.137 Inf    1   1.858  0.1264
## 
## P value adjustment: bonferroni method for 2 tests 
## Tests are performed on the log odds ratio scale
```

even though the percentage seems to be a bit different (17% vs 20%), it is not significant.

###### 2.2.1.3 Direction of rotation

the peripheral drift illusion has a rotation direction: it is perceived going counterclockwise in the current configuration. So, should be additive to a rotation direction, while substractive in the other. let’s check

```
mm_s1_r <- glmmTMB(turn~stim_type*motion*rot_direction + (1|subj), data = exp1_onlymoving, family = binomial())
simres <- simulateResiduals(mm_s1_r)
plot(simres)
```

```
## DHARMa:testOutliers with type = binomial may have inflated Type I error rates for integer-valued distributions. To get a more exact result, it is recommended to re-run testOutliers with type = 'bootstrap'. See ?testOutliers for details
```

```
Anova(mm_s1_r)
```

```
## Analysis of Deviance Table (Type II Wald chisquare tests)
## 
## Response: turn
##                                 Chisq Df Pr(>Chisq)  
## stim_type                      0.0414  1    0.83875  
## motion                         5.8227  1    0.01582 *
## rot_direction                  5.9841  1    0.01444 *
## stim_type:motion               0.4370  1    0.50859  
## stim_type:rot_direction        0.2431  1    0.62194  
## motion:rot_direction           5.0212  1    0.02504 *
## stim_type:motion:rot_direction 0.0168  1    0.89688  
## ---
## Signif. codes:  0 '***' 0.001 '**' 0.01 '*' 0.05 '.' 0.1 ' ' 1
```

There is indeed an effect of rotation direction! this is not however linked with stim type in any way, so is the same for control or for peripheral drift, so it can’t be related with the illusion. It is however dependent on the speed.

Will proceed with the post-hoc. will not include stimulus type

```
e <- emmeans(mm_s1_r, ~motion*rot_direction, type='response')
```

```
## NOTE: Results may be misleading due to involvement in interactions
```

```
e
```

```
##  motion rot_direction     prob     SE  df asymp.LCL asymp.UCL
##  2      Clockwise        0.199 0.0392 Inf     0.133     0.287
##  32     Clockwise        0.425 0.0604 Inf     0.312     0.545
##  2      CounterClockwise 0.315 0.0502 Inf     0.226     0.420
##  32     CounterClockwise 0.429 0.0602 Inf     0.317     0.549
## 
## Results are averaged over the levels of: stim_type 
## Confidence level used: 0.95 
## Intervals are back-transformed from the logit scale
```

```
contrast(e, list('CwVsCcw'=c(1/2, 1/2, -1/2, -1/2),
                 'Cw2VsCcw2'=c(1/2, 0, -1/2, 0),
                 'Cw32VsCcw32'=c(0, 1/2, 0, -1/2)),
         adjust = 'bonferroni')
```

```
##  contrast    odds.ratio     SE  df null z.ratio p.value
##  CwVsCcw          0.728 0.0968 Inf    1  -2.386  0.0512
##  Cw2VsCcw2        0.735 0.0684 Inf    1  -3.309  0.0028
##  Cw32VsCcw32      0.990 0.0941 Inf    1  -0.101  1.0000
## 
## Results are averaged over the levels of: stim_type 
## P value adjustment: bonferroni method for 3 tests 
## Tests are performed on the log odds ratio scale
```

```
# for plot

e <- emmeans(mm_s1_r, ~stim_type*motion*rot_direction, type='response')

e
```

```
##  stim_type       motion rot_direction     prob     SE  df asymp.LCL asymp.UCL
##  Control         2      Clockwise        0.186 0.0438 Inf     0.115     0.287
##  PeripheralDrift 2      Clockwise        0.212 0.0467 Inf     0.135     0.318
##  Control         32     Clockwise        0.421 0.0668 Inf     0.299     0.555
##  PeripheralDrift 32     Clockwise        0.428 0.0706 Inf     0.298     0.568
##  Control         2      CounterClockwise 0.308 0.0554 Inf     0.211     0.425
##  PeripheralDrift 2      CounterClockwise 0.322 0.0576 Inf     0.221     0.443
##  Control         32     CounterClockwise 0.447 0.0694 Inf     0.318     0.583
##  PeripheralDrift 32     CounterClockwise 0.412 0.0678 Inf     0.288     0.548
## 
## Confidence level used: 0.95 
## Intervals are back-transformed from the logit scale
```

```
write.csv(e, paste0(imagepath, 'exp1_rotDir.csv'), row.names = FALSE, dec='.')
```

```
## Warning in write.csv(e, paste0(imagepath, "exp1_rotDir.csv"), row.names =
## FALSE, : attempt to set 'dec' ignored
```

For slowly moving stimuli, response probability is much higher for counterclockwise rotating ones. I have no idea why. The absence of effect for the fast stimuli can be explained by ceiling effect: response is already at max level.

Just for completeness, I will do the contrasts also between the single simuli.

```
e
```

```
##  stim_type       motion rot_direction     prob     SE  df asymp.LCL asymp.UCL
##  Control         2      Clockwise        0.186 0.0438 Inf     0.115     0.287
##  PeripheralDrift 2      Clockwise        0.212 0.0467 Inf     0.135     0.318
##  Control         32     Clockwise        0.421 0.0668 Inf     0.299     0.555
##  PeripheralDrift 32     Clockwise        0.428 0.0706 Inf     0.298     0.568
##  Control         2      CounterClockwise 0.308 0.0554 Inf     0.211     0.425
##  PeripheralDrift 2      CounterClockwise 0.322 0.0576 Inf     0.221     0.443
##  Control         32     CounterClockwise 0.447 0.0694 Inf     0.318     0.583
##  PeripheralDrift 32     CounterClockwise 0.412 0.0678 Inf     0.288     0.548
## 
## Confidence level used: 0.95 
## Intervals are back-transformed from the logit scale
```

```
contrast(e, list('CwVsCcw'=c(1/4, 1/4, 1/4, 1/4, -1/4, -1/4, 1/4, -1/4),
                 'Illusion - CwVsCcw'=c(0, 1/2, 0, 1/2, 0, -1/2, 0, -1/2),
                 'Control - CwVsCcw'=c(1/2, 0, 1/2, 0, -1/2, 0, -1/2, 0),
                 'Illusion2 - CwVsCcw'=c(0, 1, 0, 0, 0, -1, 0, 0),
                 'Control2 - CwVsCcw'=c(1, 0, 0, 0, -1, 0, 0, 0),
                 'Illusion32 - CwVsCcw'=c(0, 0, 0, 1, 0, 0, 0, -1),
                 'Control32 - CwVsCcw'=c(0, 0, 1, 0, 0, 0, -1, 0)),
         adjust = 'bonferroni')
```

```
##  contrast             estimate    SE  df z.ratio p.value
##  CwVsCcw                -0.424 0.168 Inf  -2.519  0.0823
##  Illusion - CwVsCcw     -0.251 0.190 Inf  -1.325  1.0000
##  Control - CwVsCcw      -0.383 0.187 Inf  -2.054  0.2798
##  Illusion2 - CwVsCcw    -0.567 0.261 Inf  -2.173  0.2082
##  Control2 - CwVsCcw     -0.664 0.266 Inf  -2.497  0.0877
##  Illusion32 - CwVsCcw    0.064 0.276 Inf   0.232  1.0000
##  Control32 - CwVsCcw    -0.102 0.262 Inf  -0.391  1.0000
## 
## Note: contrasts are still on the logit scale 
## P value adjustment: bonferroni method for 7 tests
```

when splitting the data up in the two stimuli types the effects disappears

###### 2.2.1.4 Plots

###### 2.2.1.4.1 Reaction to stimuli and speeds

```
import pandas as pd
import matplotlib.pyplot as plt
imagepath = '/home/massimodeagro/Archive/Mega_sync/Experiments/Spider_PeripheralDrift/Images/'

exp1 = pd.read_csv(imagepath+'exp1.csv')
exp1Static = pd.read_csv(imagepath+'exp1_static.csv')
noneavg = exp1Static['prob'][exp1Static['stim_type'] == 'none'].item()
noneSE = exp1Static['SE'][exp1Static['stim_type'] == 'none'].item()

exp1['motion'][exp1['motion']==0] = '0'
```

```
## <string>:1: SettingWithCopyWarning: 
## A value is trying to be set on a copy of a slice from a DataFrame
## 
## See the caveats in the documentation: https://pandas.pydata.org/pandas-docs/stable/user_guide/indexing.html#returning-a-view-versus-a-copy
```

```
exp1['motion'][exp1['motion']==2] = '2'
exp1['motion'][exp1['motion']==32] = '32'

illusion = exp1[exp1['stim_type']=='PeripheralDrift']
control = exp1[exp1['stim_type']=='Control']

plt.fill_between(x=illusion['motion'], y1=noneavg - noneSE, y2=noneavg + noneSE, alpha=0.3, color='#888888')
plt.axhline(y=noneavg, color='#222222', linestyle='--')
plt.scatter(x=illusion['motion'], y=illusion['prob'], c='darkslateblue', label='Illusion')
plt.vlines(x=illusion['motion'], ymin=illusion['prob']-illusion['SE'], ymax=illusion['prob']+illusion['SE'], colors='darkslateblue')
plt.scatter(x=control['motion'], y=control['prob'], c='goldenrod', label='Control')
plt.vlines(x=control['motion'], ymin=control['prob']-control['SE'], ymax=control['prob']+control['SE'], colors='goldenrod')
plt.legend()
plt.grid(color='0.85', linestyle='--')
plt.ylim(0, 1)
```

```
## (0.0, 1.0)
```

```
plt.show()
```

###### 2.2.1.4.2 Reaction to rotation direction

```
import pandas as pd
import matplotlib.pyplot as plt
imagepath = '/home/massimodeagro/Archive/Mega_sync/Experiments/Spider_PeripheralDrift/Images/'

exp1r = pd.read_csv(imagepath+'exp1_rotDir.csv')
exp1r['motion'][exp1r['motion']==2] = '2'
```

```
## <string>:1: SettingWithCopyWarning: 
## A value is trying to be set on a copy of a slice from a DataFrame
## 
## See the caveats in the documentation: https://pandas.pydata.org/pandas-docs/stable/user_guide/indexing.html#returning-a-view-versus-a-copy
```

```
exp1r['motion'][exp1r['motion']==32] = '32'

illusion = exp1r[exp1r['stim_type']=='PeripheralDrift']
control = exp1r[exp1r['stim_type']=='Control']


illcw = control[control['rot_direction']=='Clockwise']
illccw = control[control['rot_direction']=='CounterClockwise']


plt.scatter(x=illcw['motion'], y=illcw['prob'], c='firebrick', label='Clockwise')
plt.vlines(x=illcw['motion'], ymin=illcw['prob']-illcw['SE'], ymax=illcw['prob']+illcw['SE'], colors='firebrick')
plt.scatter(x=illccw['motion'], y=illccw['prob'], c='seagreen', label='Counter-Clockwise')
plt.vlines(x=illccw['motion'], ymin=illccw['prob']-illccw['SE'], ymax=illccw['prob']+illccw['SE'], colors='seagreen')
plt.legend()
plt.grid(color='0.85', linestyle='--')
plt.ylim(0, 1)
```

```
## (0.0, 1.0)
```

```
plt.show()
```

gonna prettyfy it later

##### 2.2.2 Experiment 2 - concentric stimuli

###### 2.2.2.1 Difference between stimuli

```
mm_s2 <- glmmTMB(turn~stim_type*motion + (1|subj), data = exp2_onlystim, family = binomial())
simres <- simulateResiduals(mm_s2)
plot(simres)
```

```
Anova(mm_s2)
```

```
## Analysis of Deviance Table (Type II Wald chisquare tests)
## 
## Response: turn
##                     Chisq Df Pr(>Chisq)    
## stim_type          1.0076  1     0.3155    
## motion           156.7548  2     <2e-16 ***
## stim_type:motion   1.3783  2     0.5020    
## ---
## Signif. codes:  0 '***' 0.001 '**' 0.01 '*' 0.05 '.' 0.1 ' ' 1
```

```
e <- emmeans(mm_s2, ~stim_type*motion, type='response')

e
```

```
##  stim_type                  motion  prob     SE  df asymp.LCL asymp.UCL
##  Concentric_Control         0      0.194 0.0250 Inf     0.150     0.248
##  Concentric_PeripheralDrift 0      0.212 0.0261 Inf     0.165     0.267
##  Concentric_Control         2      0.391 0.0451 Inf     0.307     0.482
##  Concentric_PeripheralDrift 2      0.374 0.0438 Inf     0.293     0.463
##  Concentric_Control         32     0.478 0.0429 Inf     0.395     0.562
##  Concentric_PeripheralDrift 32     0.529 0.0439 Inf     0.443     0.614
## 
## Confidence level used: 0.95 
## Intervals are back-transformed from the logit scale
```

```
write.csv(e, paste0(imagepath, 'exp2.csv'), row.names = FALSE, dec='.')
```

```
## Warning in write.csv(e, paste0(imagepath, "exp2.csv"), row.names = FALSE, :
## attempt to set 'dec' ignored
```

```
contrast(e, list('speed0vs2'=c(0.5, 0.5, -0.5, -0.5, 0, 0),
                 'speed2vs32'=c(0, 0, 0.5, 0.5, -0.5, -0.5),
                 'controlVsDrift'=c(1/3, -1/3, 1/3, -1/3, 1/3, -1/3),
                 'StaticControlVsDrift'=c(1, -1, 0, 0, 0, 0),
                 'SlowControlVsDrift'=c(0, 0, 1, -1, 0, 0),
                 'FastControlVsDrift'=c(0, 0, 0, 0, 1, -1)),
         adjust = 'bonferroni')
```

```
##  contrast             odds.ratio     SE  df null z.ratio p.value
##  speed0vs2                 0.411 0.0528 Inf    1  -6.921  <.0001
##  speed2vs32                0.611 0.1061 Inf    1  -2.839  0.0272
##  controlVsDrift            0.924 0.0849 Inf    1  -0.864  1.0000
##  StaticControlVsDrift      0.900 0.1262 Inf    1  -0.754  1.0000
##  SlowControlVsDrift        1.075 0.1894 Inf    1   0.410  1.0000
##  FastControlVsDrift        0.815 0.1300 Inf    1  -1.285  1.0000
## 
## P value adjustment: bonferroni method for 6 tests 
## Tests are performed on the log odds ratio scale
```

quite exactly the same of experiment 1. Exp1 and 2 are not directly comparable, as they are different subjects, but moving stimuli are responded at a rate generally higher than experiment 1, suggesting that indeed higher moving area has an effect on detection. static stimuli are responded the same, meaning that appearance alone, independently from area, does not trigger response.

###### 2.2.2.2 Difference between static and nothing

same as above

```
mm_s2_s <- glmmTMB(turn~stim_type + (1|subj), data = exp2_onlystatic, family = binomial())
simres <- simulateResiduals(mm_s2_s)
plot(simres)
```

```
Anova(mm_s2_s)
```

```
## Analysis of Deviance Table (Type II Wald chisquare tests)
## 
## Response: turn
##            Chisq Df Pr(>Chisq)    
## stim_type 14.143  2  0.0008491 ***
## ---
## Signif. codes:  0 '***' 0.001 '**' 0.01 '*' 0.05 '.' 0.1 ' ' 1
```

```
e <- emmeans(mm_s2_s, ~stim_type, type='response')

e
```

```
##  stim_type                   prob     SE  df asymp.LCL asymp.UCL
##  Concentric_Control         0.212 0.0217 Inf     0.173     0.258
##  Concentric_PeripheralDrift 0.227 0.0222 Inf     0.186     0.273
##  none                       0.170 0.0140 Inf     0.144     0.199
## 
## Confidence level used: 0.95 
## Intervals are back-transformed from the logit scale
```

```
write.csv(e, paste0(imagepath, 'exp2_static.csv'), row.names = FALSE, dec='.')
```

```
## Warning in write.csv(e, paste0(imagepath, "exp2_static.csv"), row.names =
## FALSE, : attempt to set 'dec' ignored
```

```
contrast(e, list('ControlVsNoStim'=c(1, -1, 0),
                 'IllusionVsNoStim'=c(0, -1, 1)),
         adjust = 'bonferroni')
```

```
##  contrast         odds.ratio     SE  df null z.ratio p.value
##  ControlVsNoStim       0.918 0.1257 Inf    1  -0.628  1.0000
##  IllusionVsNoStim      0.699 0.0752 Inf    1  -3.330  0.0017
## 
## P value adjustment: bonferroni method for 2 tests 
## Tests are performed on the log odds ratio scale
```

Interestingly, here the illusory stimulus is responded at a higher rate! This is however quite peculiar. There is no statistical difference between the static control and the static illusion, and the probability of responses is similar to experiment 1. It seems that here the collected responses for the no-stimulus sections is lower. this combination creates the presence of an effect here. I believe that the preponderance of evidence here remains in the direction of “no illusory perception”, but let’s keep our mind open.

###### 2.2.2.3 Direction of rotation

```
mm_s2_r <- glmmTMB(turn~stim_type*motion*rot_direction + (1|subj), data = exp2_onlymoving, family = binomial())
simres <- simulateResiduals(mm_s2_r)
plot(simres)
```

```
Anova(mm_s2_r)
```

```
## Analysis of Deviance Table (Type II Wald chisquare tests)
## 
## Response: turn
##                                  Chisq Df Pr(>Chisq)    
## stim_type                       0.7562  1  0.3845132    
## motion                          0.0419  1  0.8378169    
## rot_direction                   1.3532  1  0.2447208    
## stim_type:motion                1.3848  1  0.2392828    
## stim_type:rot_direction        14.2290  1  0.0001619 ***
## motion:rot_direction            0.9837  1  0.3212800    
## stim_type:motion:rot_direction  6.9545  1  0.0083611 ** 
## ---
## Signif. codes:  0 '***' 0.001 '**' 0.01 '*' 0.05 '.' 0.1 ' ' 1
```

In this case, the stimulus type is relevant! Quite peculiar. Let’s check

```
e <- emmeans(mm_s2_r, ~stim_type*motion*rot_direction, type='response')

e
```

```
##  stim_type                  motion rot_direction     prob     SE  df asymp.LCL
##  Concentric_Control         2      Clockwise        0.513 0.0670 Inf     0.384
##  Concentric_PeripheralDrift 2      Clockwise        0.303 0.0578 Inf     0.203
##  Concentric_Control         32     Clockwise        0.448 0.0565 Inf     0.341
##  Concentric_PeripheralDrift 32     Clockwise        0.459 0.0593 Inf     0.347
##  Concentric_Control         2      CounterClockwise 0.381 0.0636 Inf     0.266
##  Concentric_PeripheralDrift 2      CounterClockwise 0.570 0.0645 Inf     0.442
##  Concentric_Control         32     CounterClockwise 0.412 0.0582 Inf     0.304
##  Concentric_PeripheralDrift 32     CounterClockwise 0.512 0.0598 Inf     0.397
##  asymp.UCL
##      0.641
##      0.427
##      0.559
##      0.576
##      0.511
##      0.690
##      0.529
##      0.627
## 
## Confidence level used: 0.95 
## Intervals are back-transformed from the logit scale
```

```
write.csv(e, paste0(imagepath, 'exp2_rotDir.csv'), row.names = FALSE, dec='.')
```

```
## Warning in write.csv(e, paste0(imagepath, "exp2_rotDir.csv"), row.names =
## FALSE, : attempt to set 'dec' ignored
```

```
contrast(e, list('CwVsCcw'=c(1/4, 1/4, 1/4, 1/4, -1/4, -1/4, 1/4, -1/4),
                 'Illusion - CwVsCcw'=c(0, 1/2, 0, 1/2, 0, -1/2, 0, -1/2),
                 'Control - CwVsCcw'=c(1/2, 0, 1/2, 0, -1/2, 0, -1/2, 0),
                 'Illusion2 - CwVsCcw'=c(0, 1, 0, 0, 0, -1, 0, 0),
                 'Control2 - CwVsCcw'=c(1, 0, 0, 0, -1, 0, 0, 0),
                 'Illusion32 - CwVsCcw'=c(0, 0, 0, 1, 0, 0, 0, -1),
                 'Control32 - CwVsCcw'=c(0, 0, 1, 0, 0, 0, -1, 0)),
         adjust = 'bonferroni')
```

```
##  contrast             estimate    SE  df z.ratio p.value
##  CwVsCcw                -0.338 0.150 Inf  -2.253  0.1700
##  Illusion - CwVsCcw     -0.663 0.175 Inf  -3.788  0.0011
##  Control - CwVsCcw       0.342 0.172 Inf   1.991  0.3253
##  Illusion2 - CwVsCcw    -1.114 0.260 Inf  -4.285  0.0001
##  Control2 - CwVsCcw      0.539 0.262 Inf   2.060  0.2757
##  Illusion32 - CwVsCcw   -0.213 0.234 Inf  -0.907  1.0000
##  Control32 - CwVsCcw     0.146 0.223 Inf   0.653  1.0000
## 
## Note: contrasts are still on the logit scale 
## P value adjustment: bonferroni method for 7 tests
```

there is a preference for counterclockwise DRIVEN BY illusion at slow speed. Also is in the wrong direction! I need a new experiment, testing what happens by flipping the illusion stimulus.

###### 2.2.2.4 Plots

###### 2.2.2.4.1 Reaction to stimuli and speeds

```
import matplotlib.pyplot as plt
import pandas as pd
imagepath = '/home/massimodeagro/Archive/Mega_sync/Experiments/Spider_PeripheralDrift/Images/'

exp2 = pd.read_csv(imagepath+'exp2.csv')
exp2Static = pd.read_csv(imagepath+'exp2_static.csv')
noneavg = exp2Static['prob'][exp2Static['stim_type'] == 'none'].item()
noneSE = exp2Static['SE'][exp2Static['stim_type'] == 'none'].item()

exp2['motion'][exp2['motion']==0] = '0'
```

```
## <string>:1: SettingWithCopyWarning: 
## A value is trying to be set on a copy of a slice from a DataFrame
## 
## See the caveats in the documentation: https://pandas.pydata.org/pandas-docs/stable/user_guide/indexing.html#returning-a-view-versus-a-copy
```

```
exp2['motion'][exp2['motion']==2] = '2'
exp2['motion'][exp2['motion']==32] = '32'

illusion = exp2[exp2['stim_type']=='Concentric_PeripheralDrift']
control = exp2[exp2['stim_type']=='Concentric_Control']

plt.fill_between(x=illusion['motion'], y1=noneavg - noneSE, y2=noneavg + noneSE, alpha=0.3, color='#888888')
plt.axhline(y=noneavg, color='#222222', linestyle='--')
plt.scatter(x=illusion['motion'], y=illusion['prob'], c='darkslateblue', label='Illusion')
plt.vlines(x=illusion['motion'], ymin=illusion['prob']-illusion['SE'], ymax=illusion['prob']+illusion['SE'], colors='darkslateblue')
plt.scatter(x=control['motion'], y=control['prob'], c='goldenrod', label='Control')
plt.vlines(x=control['motion'], ymin=control['prob']-control['SE'], ymax=control['prob']+control['SE'], colors='goldenrod')
plt.legend()
plt.grid(color='0.85', linestyle='--')
plt.ylim(0, 1)
```

```
## (0.0, 1.0)
```

```
plt.show()
```

###### 2.2.2.4.2 Reaction to rotation direction

```
import pandas as pd
import matplotlib.pyplot as plt
imagepath = '/home/massimodeagro/Archive/Mega_sync/Experiments/Spider_PeripheralDrift/Images/'

exp2r = pd.read_csv(imagepath+'exp2_rotDir.csv')
exp2r['motion'][exp2r['motion']==2] = '2'
```

```
## <string>:1: SettingWithCopyWarning: 
## A value is trying to be set on a copy of a slice from a DataFrame
## 
## See the caveats in the documentation: https://pandas.pydata.org/pandas-docs/stable/user_guide/indexing.html#returning-a-view-versus-a-copy
```

```
exp2r['motion'][exp2r['motion']==32] = '32'

slow = exp2r[exp2r['motion']=='2']
fast = exp2r[exp2r['motion']=='32']

illcw = slow[slow['rot_direction']=='Clockwise']
illccw = slow[slow['rot_direction']=='CounterClockwise']

plt.scatter(x=illcw['stim_type'], y=illcw['prob'], c='firebrick', label='Clockwise')
plt.vlines(x=illcw['stim_type'], ymin=illcw['prob']-illcw['SE'], ymax=illcw['prob']+illcw['SE'], colors='firebrick')
plt.scatter(x=illccw['stim_type'], y=illccw['prob'], c='seagreen', label='Counter-Clockwise')
plt.vlines(x=illccw['stim_type'], ymin=illccw['prob']-illccw['SE'], ymax=illccw['prob']+illccw['SE'], colors='seagreen')
plt.legend()
plt.grid(color='0.85', linestyle='--')
plt.ylim(0, 1)
```

```
## (0.0, 1.0)
```

```
plt.show()
```

gonna prettyfy it later

##### 2.2.3 Experiment 3 - flipped concentric stimuli

###### 2.2.3.1 Difference between stimuli

```
mm_s3 <- glmmTMB(turn~stim_type*motion + (1|subj), data = exp3_onlystim, family = binomial())
simres <- simulateResiduals(mm_s3)
plot(simres)
```

```
Anova(mm_s3)
```

```
## Analysis of Deviance Table (Type II Wald chisquare tests)
## 
## Response: turn
##                    Chisq Df Pr(>Chisq)    
## stim_type         1.0004  1     0.3172    
## motion           17.7230  1  2.555e-05 ***
## stim_type:motion  1.5297  1     0.2162    
## ---
## Signif. codes:  0 '***' 0.001 '**' 0.01 '*' 0.05 '.' 0.1 ' ' 1
```

```
e <- emmeans(mm_s3, ~stim_type*motion, type='response')

e
```

```
##  stim_type                          motion  prob     SE  df asymp.LCL asymp.UCL
##  Concentric_Control                 0      0.194 0.0328 Inf     0.138     0.267
##  Concentric_PeripheralDrift_flipped 0      0.186 0.0314 Inf     0.133     0.256
##  Concentric_Control                 2      0.261 0.0390 Inf     0.192     0.345
##  Concentric_PeripheralDrift_flipped 2      0.318 0.0423 Inf     0.242     0.406
## 
## Confidence level used: 0.95 
## Intervals are back-transformed from the logit scale
```

```
write.csv(e, paste0(imagepath, 'exp3.csv'), row.names = FALSE, dec='.')
```

```
## Warning in write.csv(e, paste0(imagepath, "exp3.csv"), row.names = FALSE, :
## attempt to set 'dec' ignored
```

```
contrast(e, list('speed0vs2'=c(0.5, 0.5, -0.5, -0.5),
                 'StaticControlVsDrift'=c(0, 0, 1, -1)),
         adjust = 'bonferroni')
```

```
##  contrast             odds.ratio     SE  df null z.ratio p.value
##  speed0vs2                 0.578 0.0769 Inf    1  -4.118  0.0001
##  StaticControlVsDrift      0.757 0.1342 Inf    1  -1.570  0.2328
## 
## P value adjustment: bonferroni method for 2 tests 
## Tests are performed on the log odds ratio scale
```

As it happened until now, no difference whatsoever between control and illusion. let’s proceed

###### 2.2.3.2 Difference between static and nothing

same as above

```
mm_s3_s <- glmmTMB(turn~stim_type + (1|subj), data = exp3_onlystatic, family = binomial())
simres <- simulateResiduals(mm_s3_s)
plot(simres)
```

```
Anova(mm_s3_s)
```

```
## Analysis of Deviance Table (Type II Wald chisquare tests)
## 
## Response: turn
##            Chisq Df Pr(>Chisq)  
## stim_type 5.2094  2    0.07393 .
## ---
## Signif. codes:  0 '***' 0.001 '**' 0.01 '*' 0.05 '.' 0.1 ' ' 1
```

```
e <- emmeans(mm_s3_s, ~stim_type, type='response')

e
```

```
##  stim_type                           prob     SE  df asymp.LCL asymp.UCL
##  Concentric_Control                 0.208 0.0276 Inf     0.159     0.267
##  Concentric_PeripheralDrift_flipped 0.197 0.0260 Inf     0.151     0.253
##  none                               0.162 0.0163 Inf     0.132     0.196
## 
## Confidence level used: 0.95 
## Intervals are back-transformed from the logit scale
```

```
write.csv(e, paste0(imagepath, 'exp3_static.csv'), row.names = FALSE, dec='.')
```

```
## Warning in write.csv(e, paste0(imagepath, "exp3_static.csv"), row.names =
## FALSE, : attempt to set 'dec' ignored
```

```
contrast(e, list('ControlVsNoStim'=c(1, 0, -1),
                 'IllusionVsNoStim'=c(0, 1, -1)),
         adjust = 'bonferroni')
```

```
##  contrast         odds.ratio    SE  df null z.ratio p.value
##  ControlVsNoStim        1.36 0.215 Inf    1   1.951  0.1022
##  IllusionVsNoStim       1.28 0.196 Inf    1   1.592  0.2228
## 
## P value adjustment: bonferroni method for 2 tests 
## Tests are performed on the log odds ratio scale
```

No difference. again I feel confident in confirming that appearing stimuli are not enough to trigger a response

###### 2.2.3.3 Direction of rotation

```
mm_s3_r <- glmmTMB(turn~stim_type*rot_direction + (1|subj), data = exp3_onlymoving, family = binomial())
simres <- simulateResiduals(mm_s3_r)
plot(simres)
```

```
Anova(mm_s3_r)
```

```
## Analysis of Deviance Table (Type II Wald chisquare tests)
## 
## Response: turn
##                           Chisq Df Pr(>Chisq)    
## stim_type                2.4979  1  0.1139985    
## rot_direction           11.5823  1  0.0006658 ***
## stim_type:rot_direction  8.2039  1  0.0041800 ** 
## ---
## Signif. codes:  0 '***' 0.001 '**' 0.01 '*' 0.05 '.' 0.1 ' ' 1
```

ah ah! same effect as before. let’s inquire further

```
e <- emmeans(mm_s3_r, ~stim_type*rot_direction, type='response')

e
```

```
##  stim_type                          rot_direction     prob     SE  df asymp.LCL
##  Concentric_Control                 Clockwise        0.268 0.0480 Inf     0.184
##  Concentric_PeripheralDrift_flipped Clockwise        0.434 0.0575 Inf     0.327
##  Concentric_Control                 CounterClockwise 0.251 0.0487 Inf     0.168
##  Concentric_PeripheralDrift_flipped CounterClockwise 0.196 0.0410 Inf     0.128
##  asymp.UCL
##      0.371
##      0.548
##      0.358
##      0.289
## 
## Confidence level used: 0.95 
## Intervals are back-transformed from the logit scale
```

```
write.csv(e, paste0(imagepath, 'exp3_rotDir.csv'), row.names = FALSE, dec='.')
```

```
## Warning in write.csv(e, paste0(imagepath, "exp3_rotDir.csv"), row.names =
## FALSE, : attempt to set 'dec' ignored
```

```
contrast(e, list('CwVsCcw'=c(1/2, 1/2, -1/2, -1/2),
                 'Illusion - CwVsCcw'=c(0, 1, 0, -1),
                 'Control - CwVsCcw'=c(1, 0, -1, 0)),
         adjust = 'bonferroni')
```

```
##  contrast           odds.ratio    SE  df null z.ratio p.value
##  CwVsCcw                  1.85 0.344 Inf    1   3.304  0.0029
##  Illusion / CwVsCcw       3.14 0.809 Inf    1   4.438  <.0001
##  Control / CwVsCcw        1.09 0.290 Inf    1   0.316  1.0000
## 
## P value adjustment: bonferroni method for 3 tests 
## Tests are performed on the log odds ratio scale
```

Indeed, here is the same effect as experiment 2, but flipped as the stimuli are

###### 2.2.3.4 Plots

###### 2.2.3.4.1 Reaction to stimuli and speeds

```
import matplotlib.pyplot as plt
import pandas as pd
imagepath = '/home/massimodeagro/Archive/Mega_sync/Experiments/Spider_PeripheralDrift/Images/'

exp3 = pd.read_csv(imagepath+'exp3.csv')
exp3Static = pd.read_csv(imagepath+'exp3_static.csv')
noneavg = exp3Static['prob'][exp3Static['stim_type'] == 'none'].item()
noneSE = exp3Static['SE'][exp3Static['stim_type'] == 'none'].item()

exp3['motion'][exp3['motion']==0] = '0'
```

```
## <string>:1: SettingWithCopyWarning: 
## A value is trying to be set on a copy of a slice from a DataFrame
## 
## See the caveats in the documentation: https://pandas.pydata.org/pandas-docs/stable/user_guide/indexing.html#returning-a-view-versus-a-copy
```

```
exp3['motion'][exp3['motion']==2] = '2'

illusion = exp3[exp3['stim_type']=='Concentric_PeripheralDrift_flipped']
control = exp3[exp3['stim_type']=='Concentric_Control']

plt.fill_between(x=illusion['motion'], y1=noneavg - noneSE, y2=noneavg + noneSE, alpha=0.3, color='#888888')
plt.axhline(y=noneavg, color='#222222', linestyle='--')
plt.scatter(x=illusion['motion'], y=illusion['prob'], c='darkslateblue', label='Illusion')
plt.vlines(x=illusion['motion'], ymin=illusion['prob']-illusion['SE'], ymax=illusion['prob']+illusion['SE'], colors='darkslateblue')
plt.scatter(x=control['motion'], y=control['prob'], c='goldenrod', label='Control')
plt.vlines(x=control['motion'], ymin=control['prob']-control['SE'], ymax=control['prob']+control['SE'], colors='goldenrod')
plt.legend()
plt.grid(color='0.85', linestyle='--')
plt.ylim(0, 1)
```

```
## (0.0, 1.0)
```

```
plt.show()
```

###### 2.2.3.4.2 Reaction to rotation direction

```
import pandas as pd
import matplotlib.pyplot as plt
imagepath = '/home/massimodeagro/Archive/Mega_sync/Experiments/Spider_PeripheralDrift/Images/'

exp3r = pd.read_csv(imagepath+'exp3_rotDir.csv')

illcw = exp3r[exp3r['rot_direction']=='Clockwise']
illccw = exp3r[exp3r['rot_direction']=='CounterClockwise']

plt.scatter(x=illcw['stim_type'], y=illcw['prob'], c='firebrick', label='Clockwise')
plt.vlines(x=illcw['stim_type'], ymin=illcw['prob']-illcw['SE'], ymax=illcw['prob']+illcw['SE'], colors='firebrick')
plt.scatter(x=illccw['stim_type'], y=illccw['prob'], c='seagreen', label='Counter-Clockwise')
plt.vlines(x=illccw['stim_type'], ymin=illccw['prob']-illccw['SE'], ymax=illccw['prob']+illccw['SE'], colors='seagreen')
plt.legend()
plt.grid(color='0.85', linestyle='--')
plt.ylim(0, 1)
```

```
## (0.0, 1.0)
```

```
plt.show()
```
