## Supplementary figures and images for "Jumping spiders are not fooled by the peripheral drift illusion"

### unnamed-chunk-7-1.png

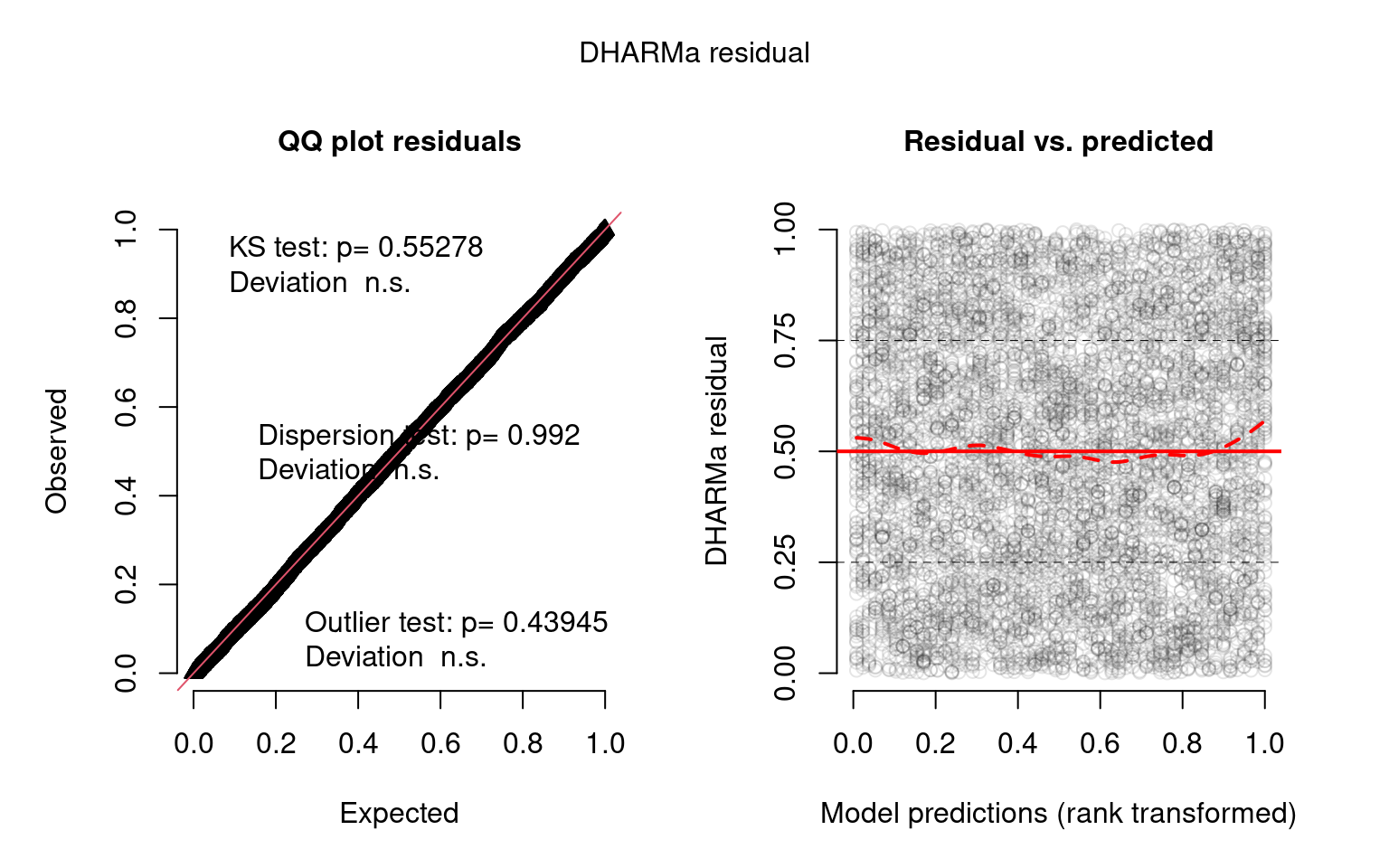

### unnamed-chunk-8-1.png

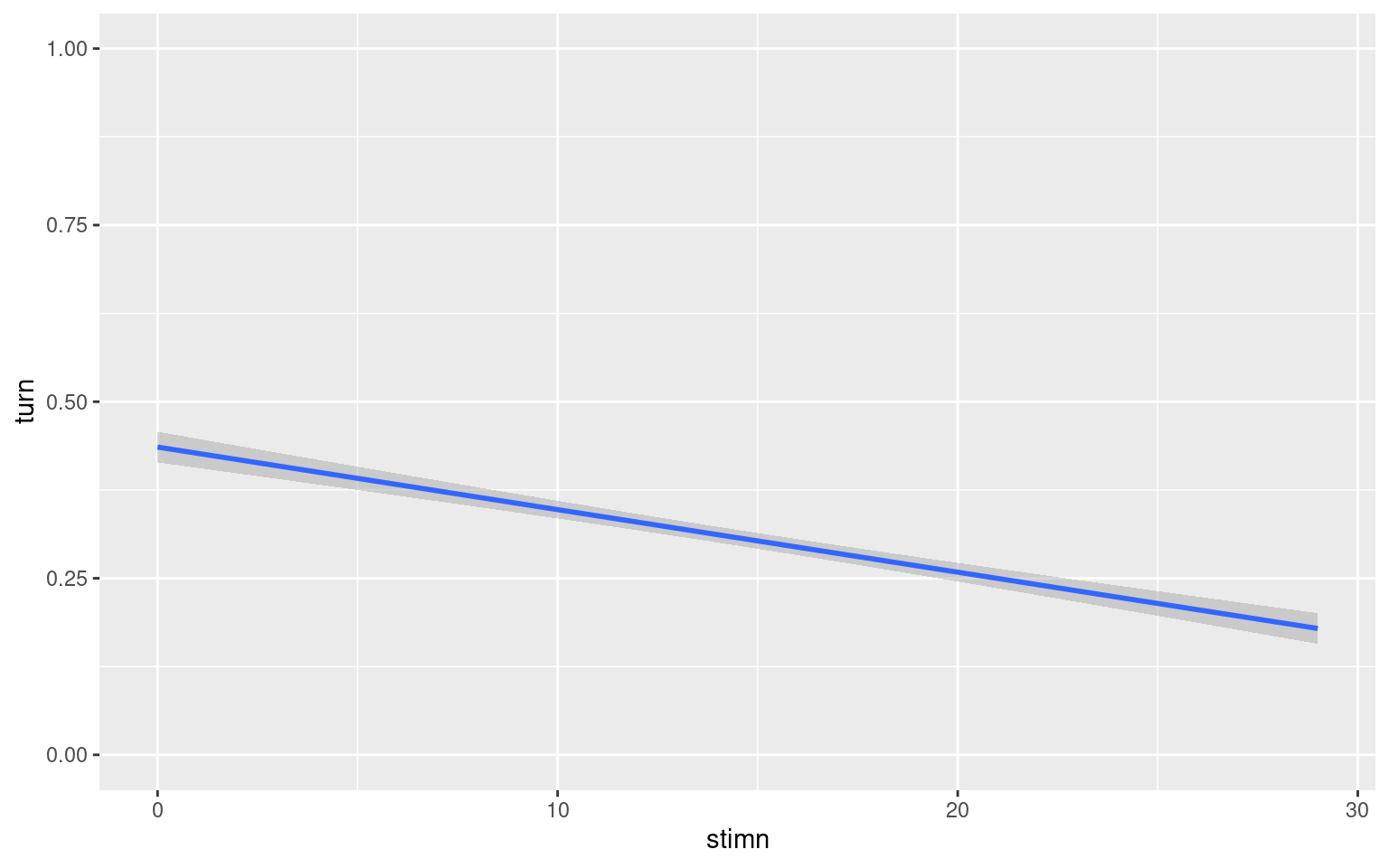

### unnamed-chunk-10-1.png

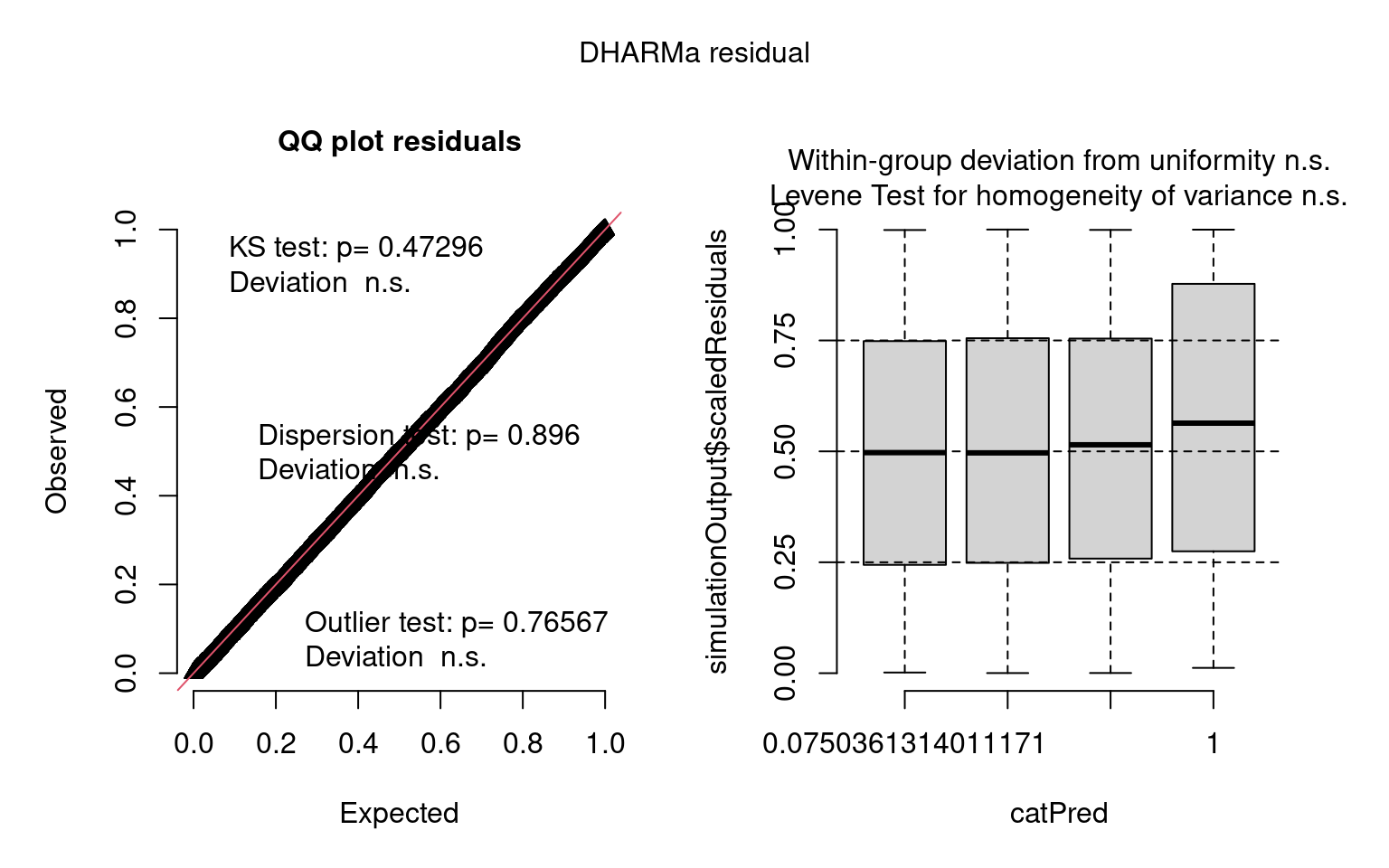

### unnamed-chunk-11-1.png

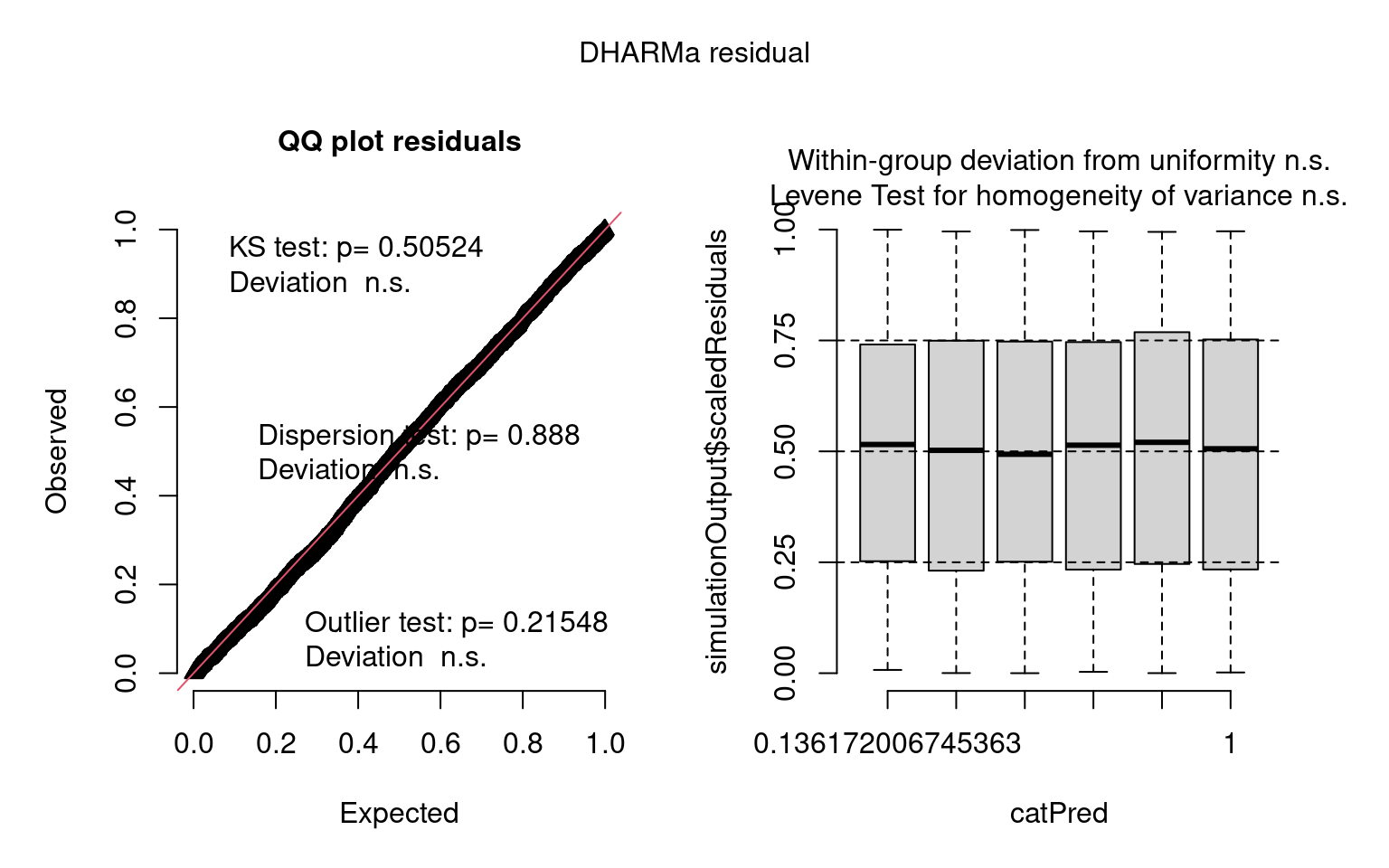

### unnamed-chunk-12-1.png

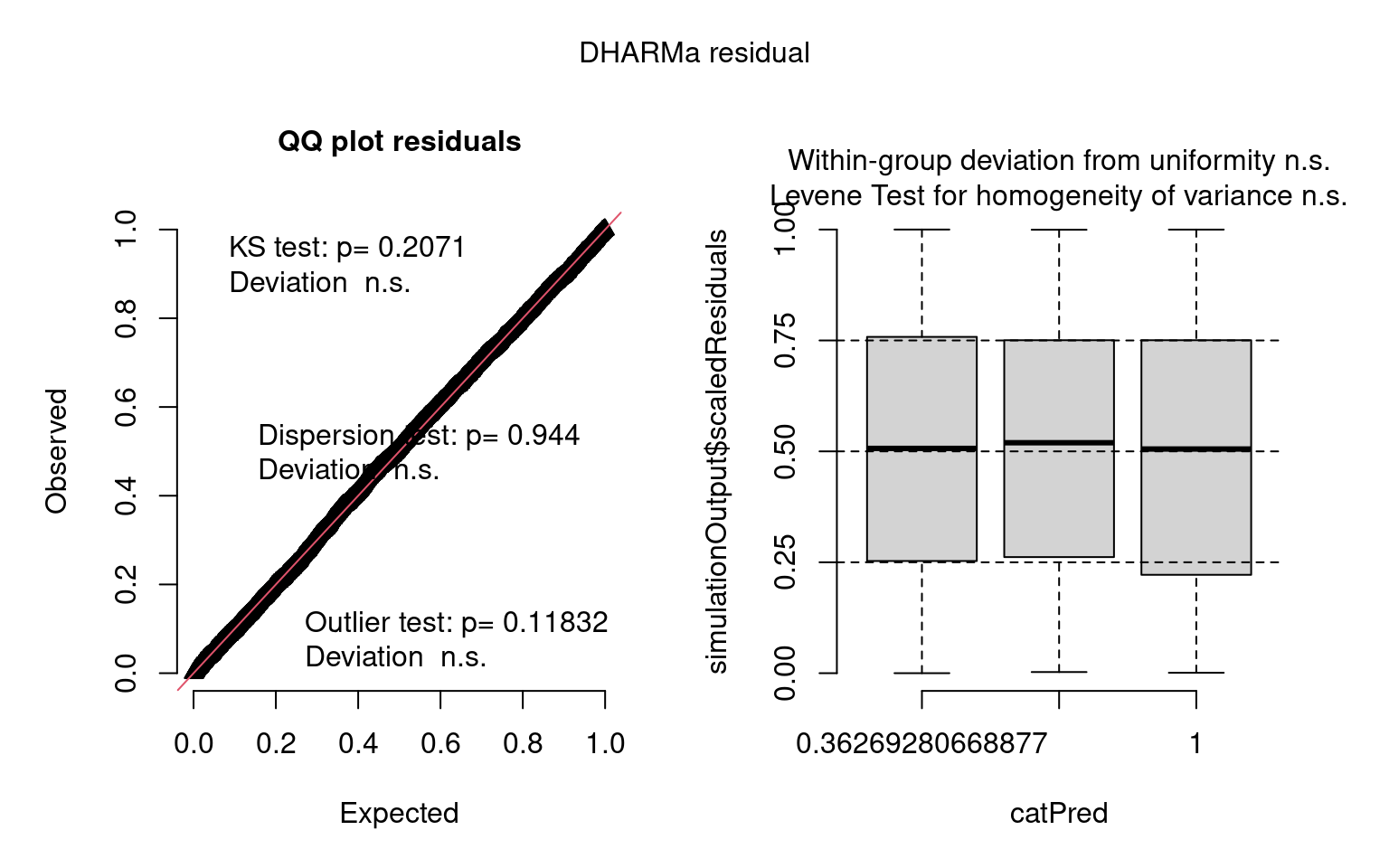

### unnamed-chunk-13-1.png

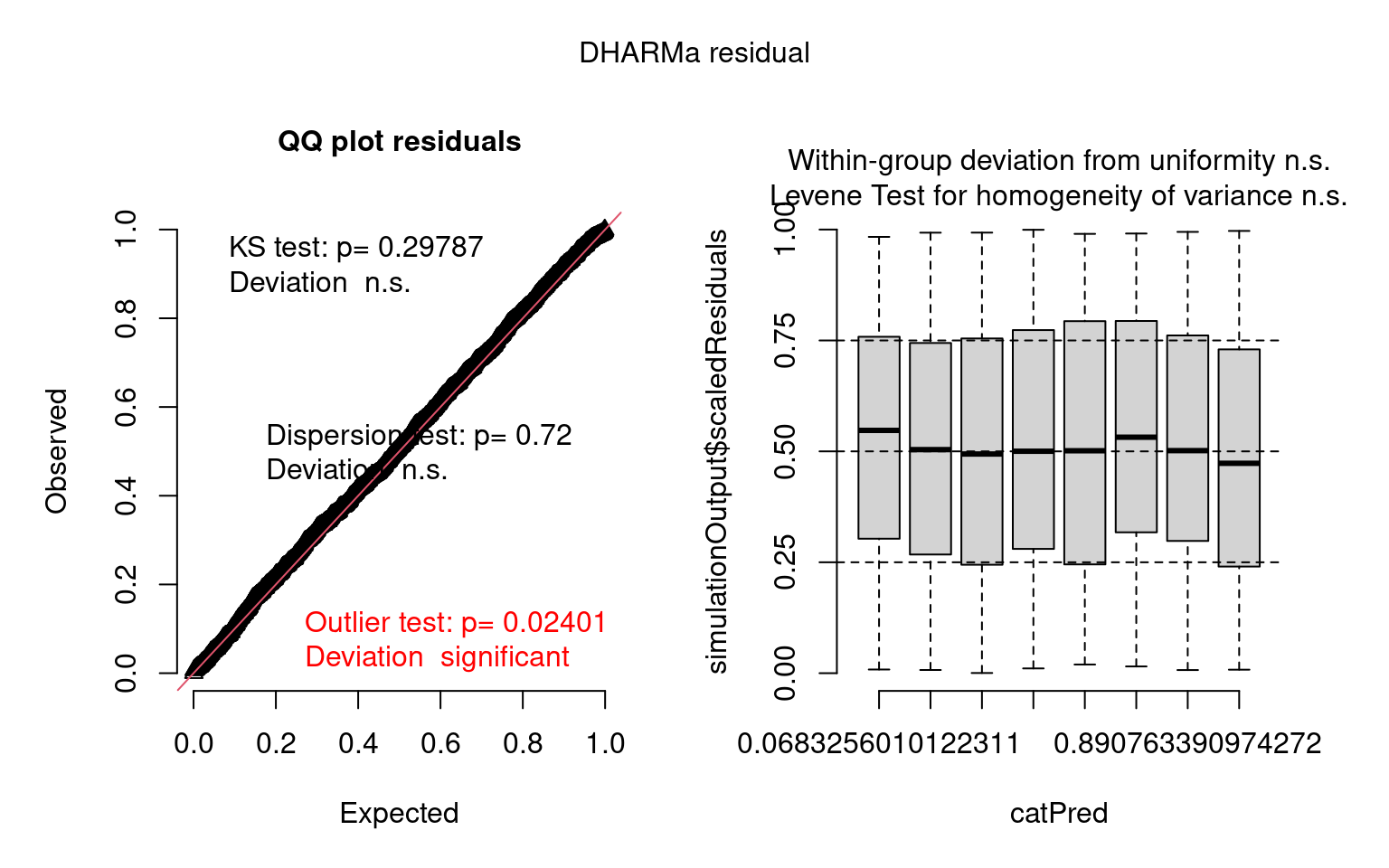

### unnamed-chunk-16-1.png

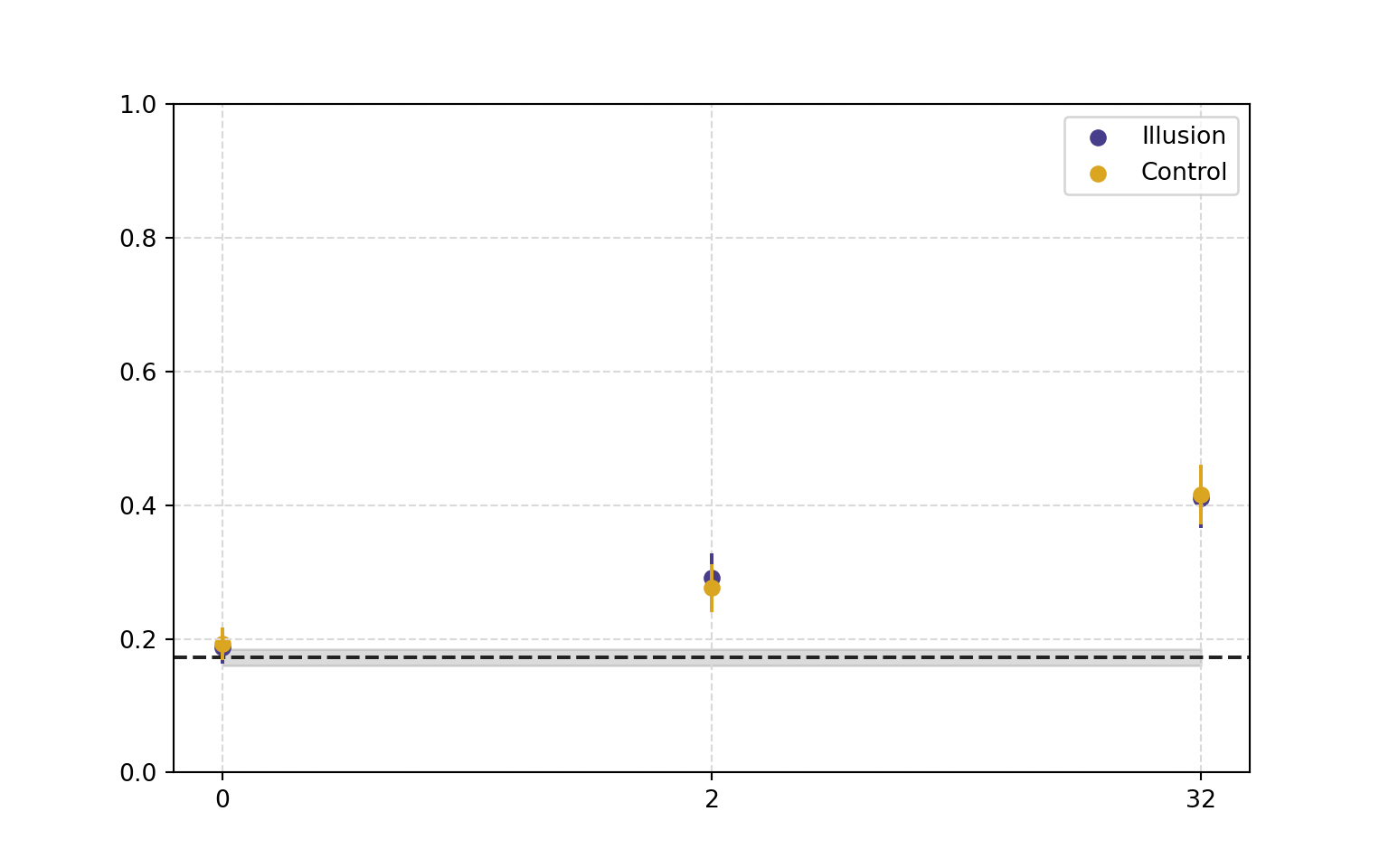

### unnamed-chunk-17-3.png

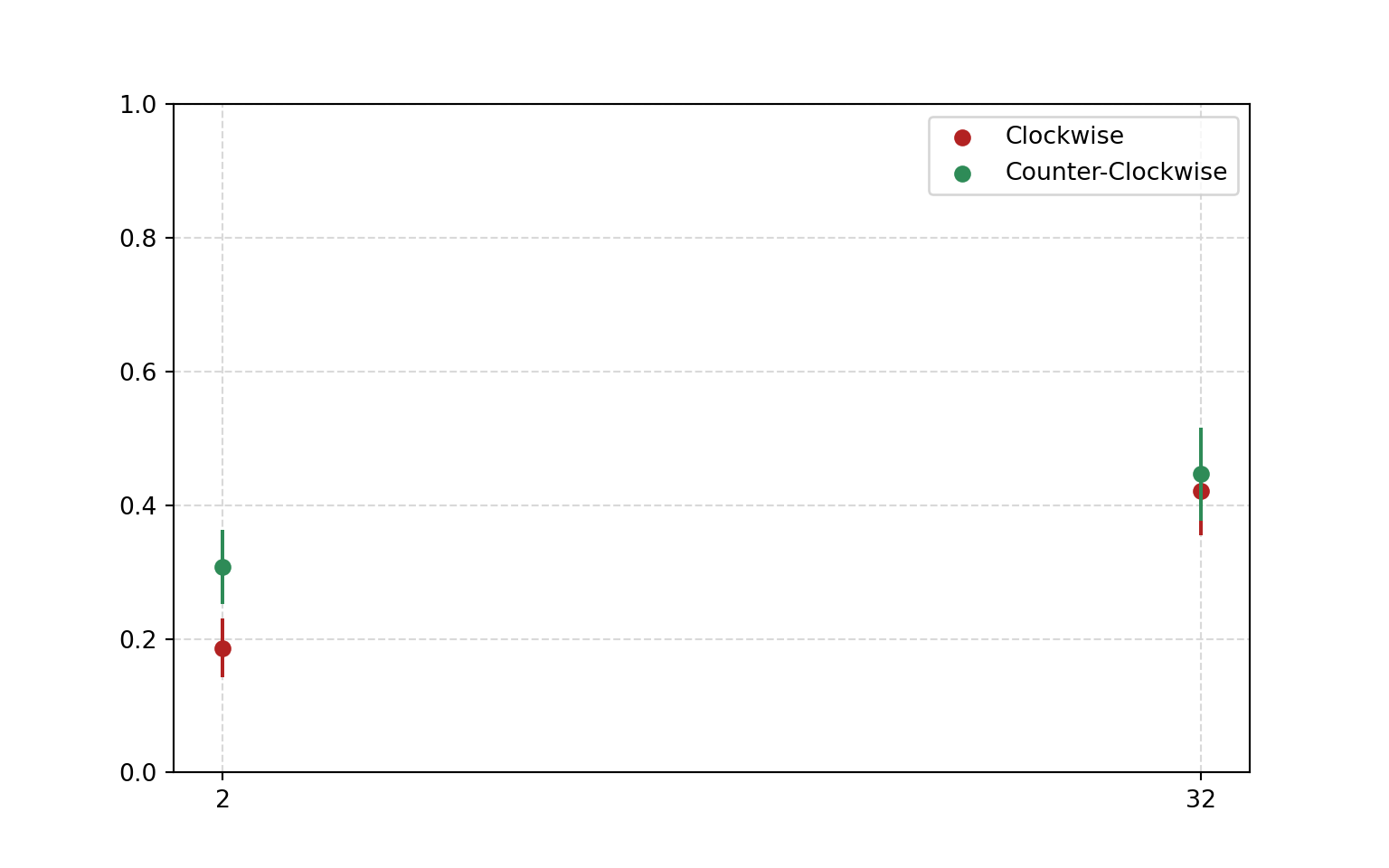

### unnamed-chunk-18-5.png

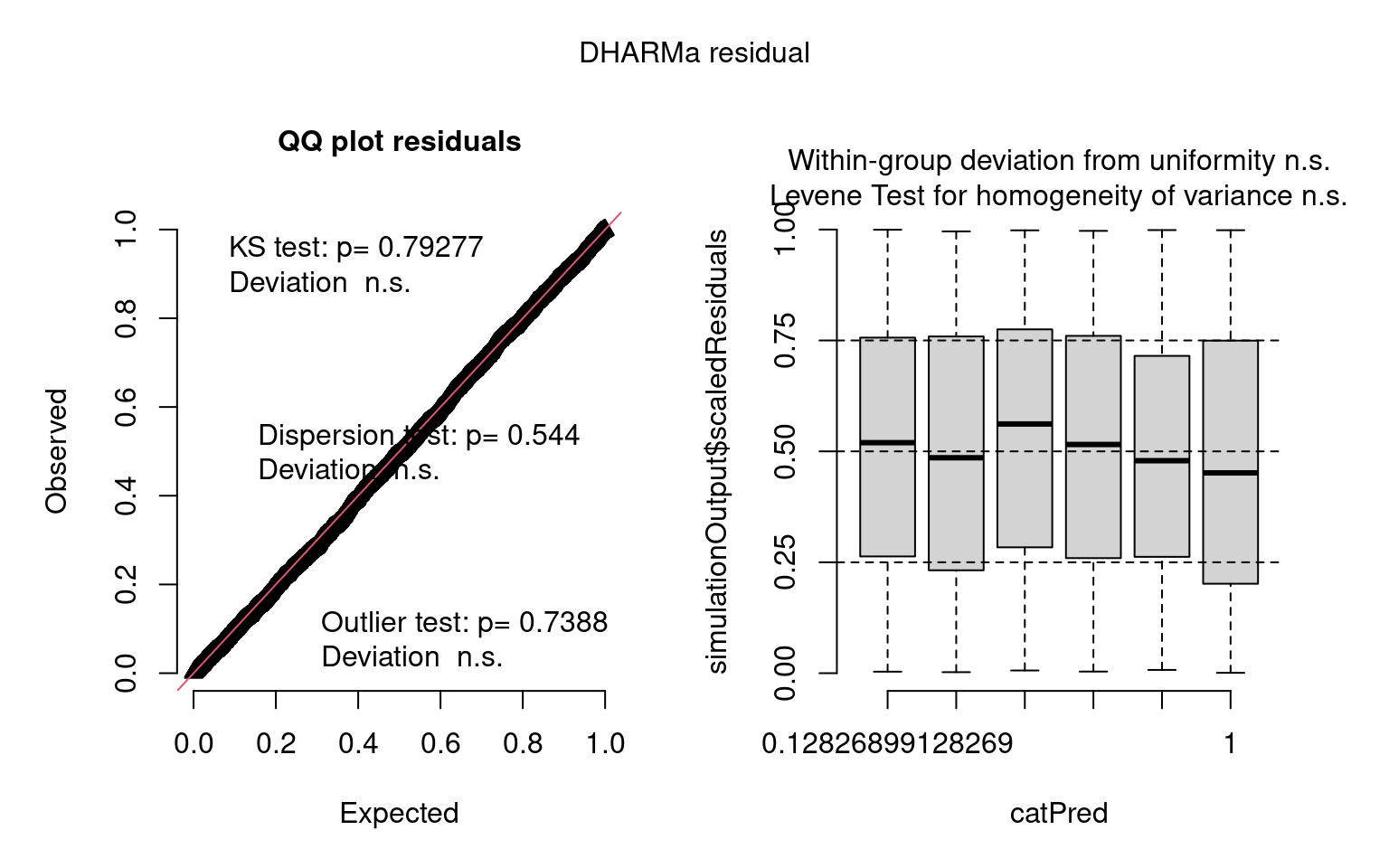

### unnamed-chunk-19-1.png

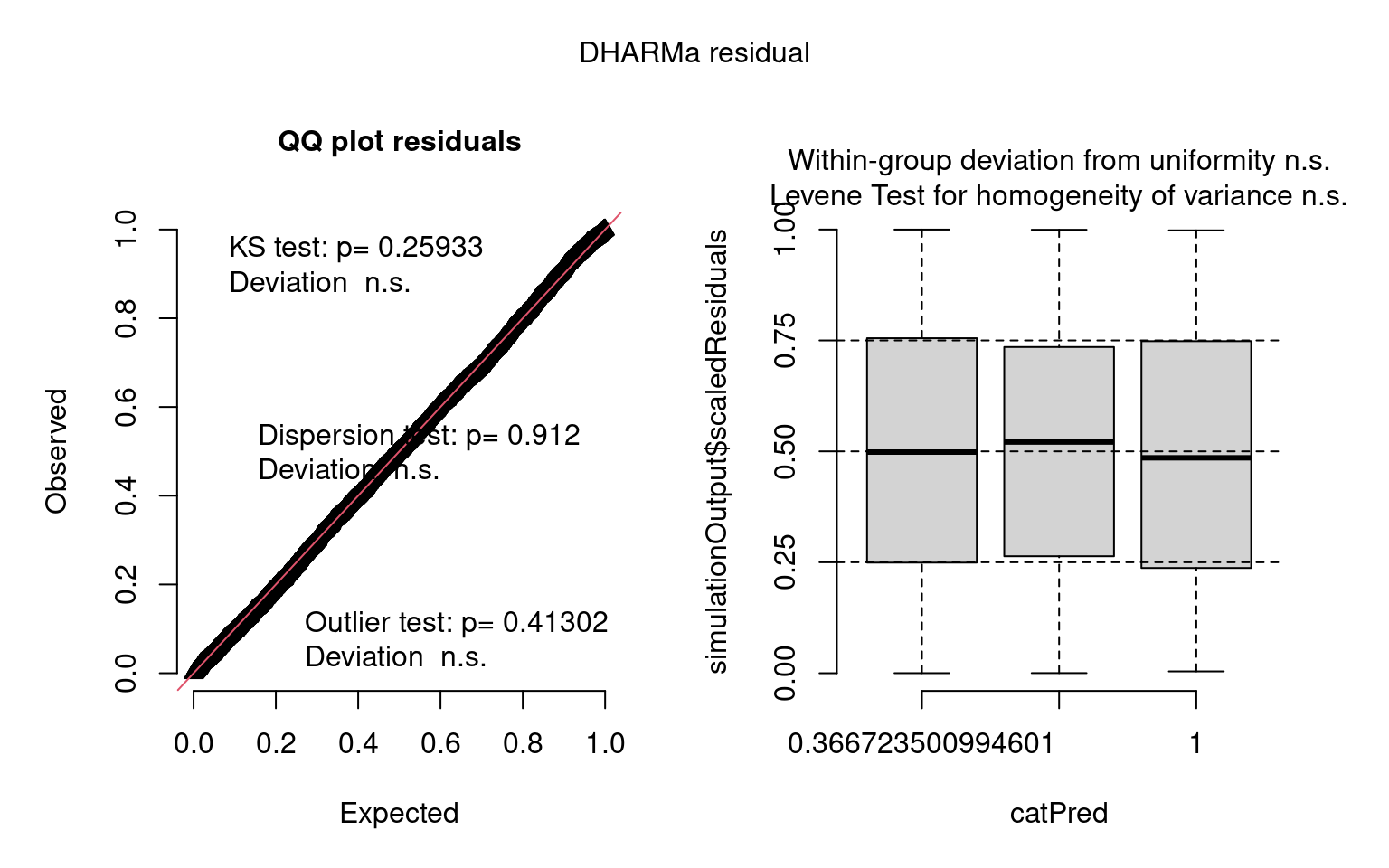

### unnamed-chunk-20-1.png

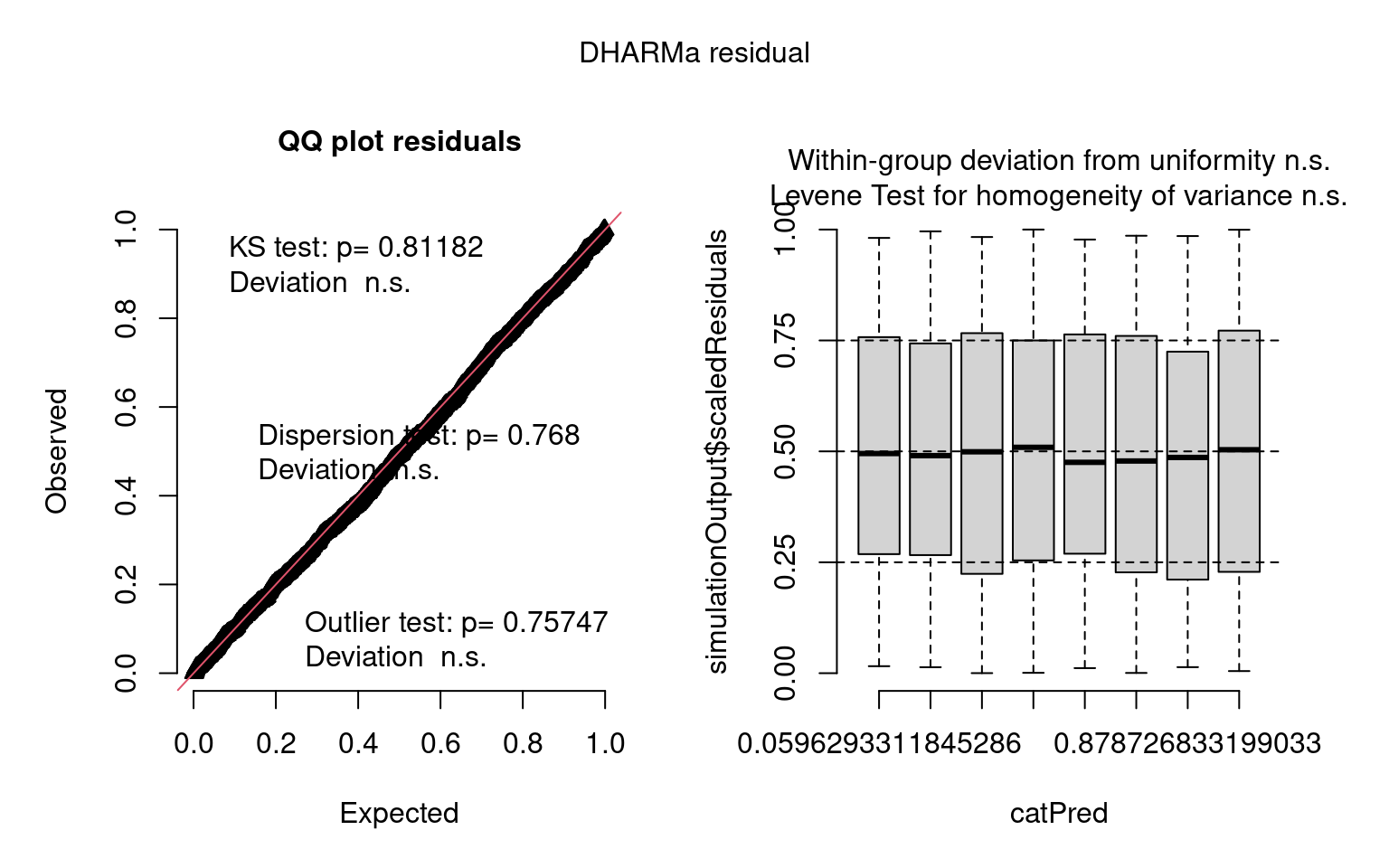

### unnamed-chunk-22-1.png

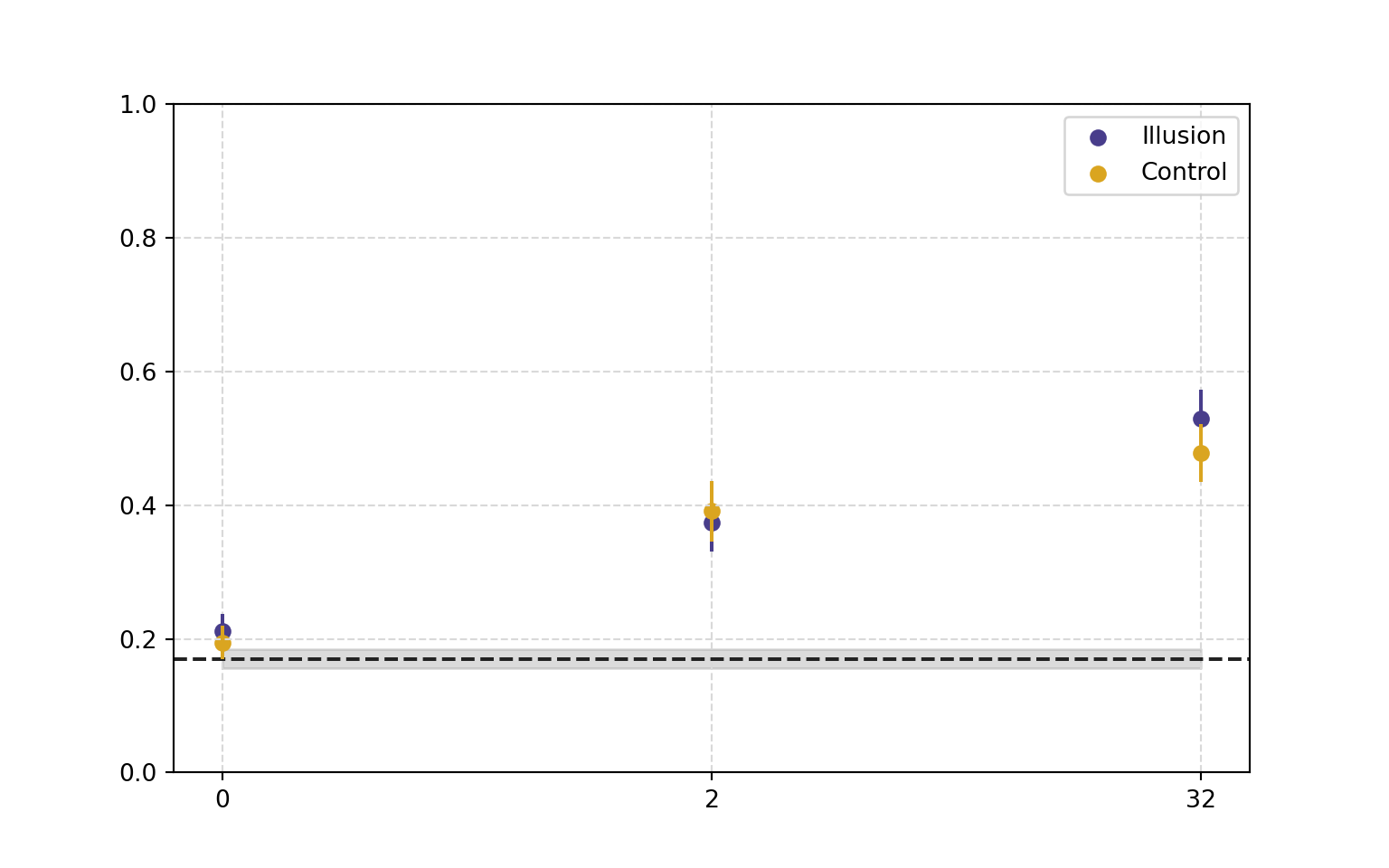

### unnamed-chunk-23-3.png

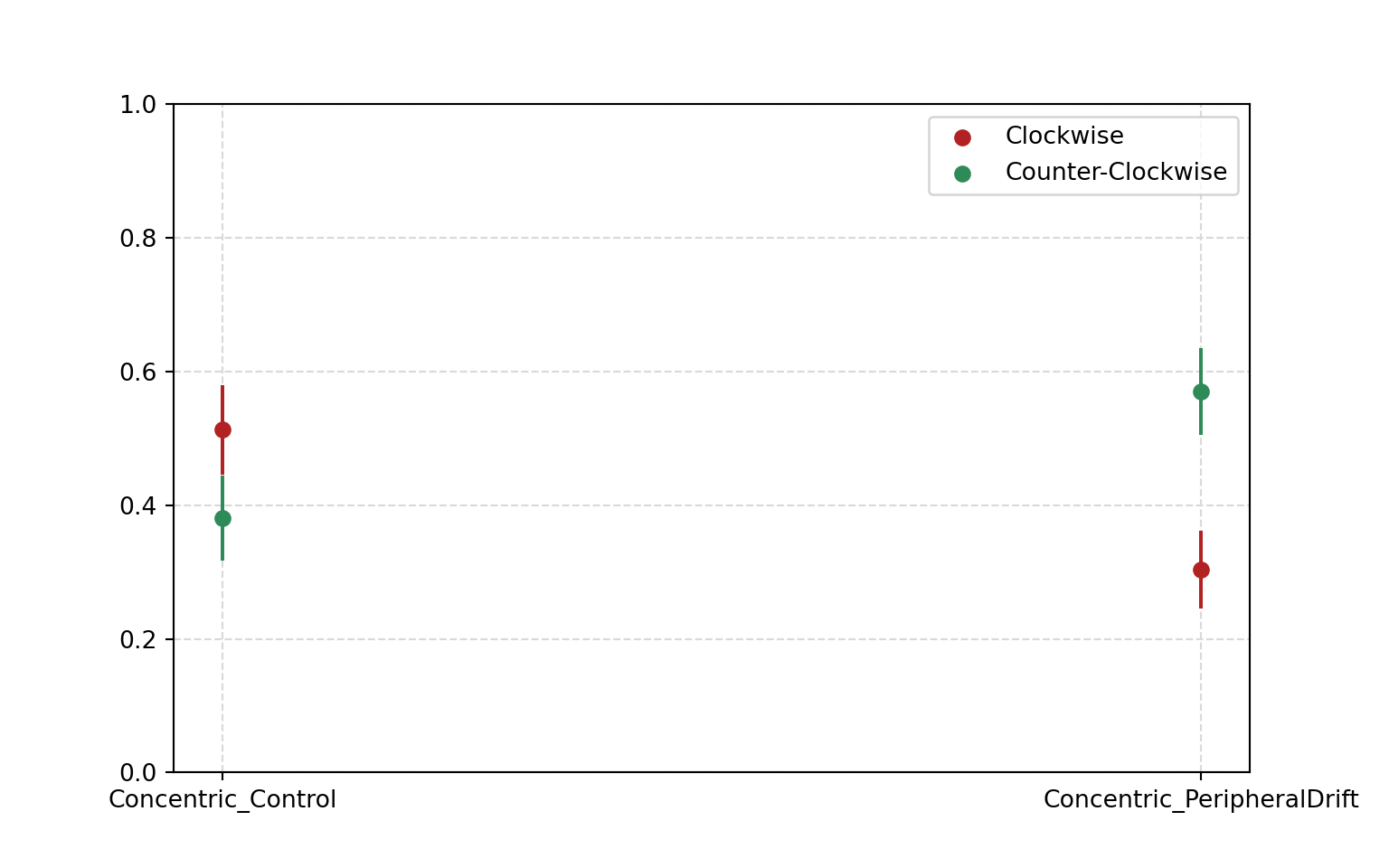

### unnamed-chunk-24-5.png

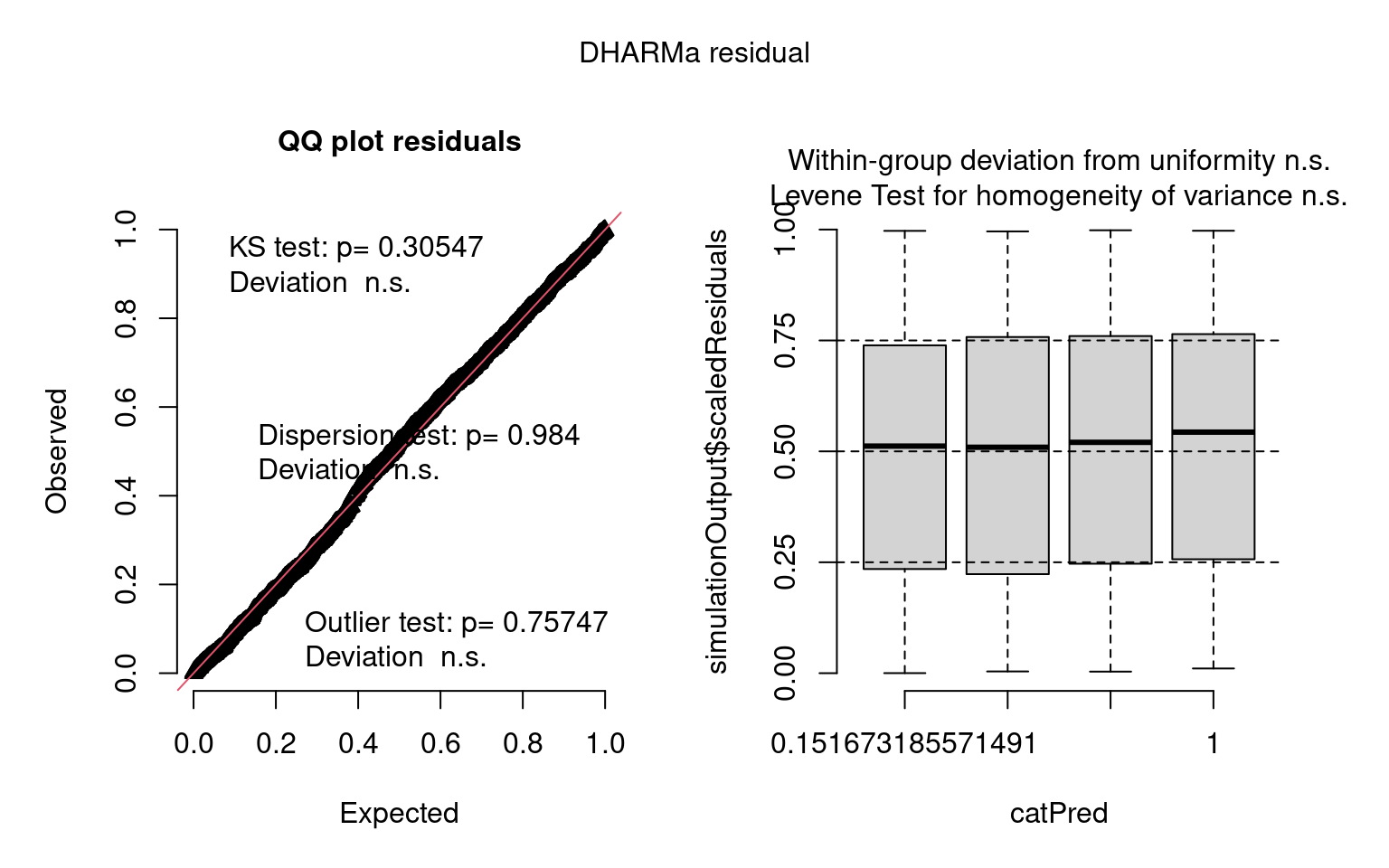

### unnamed-chunk-25-1.png

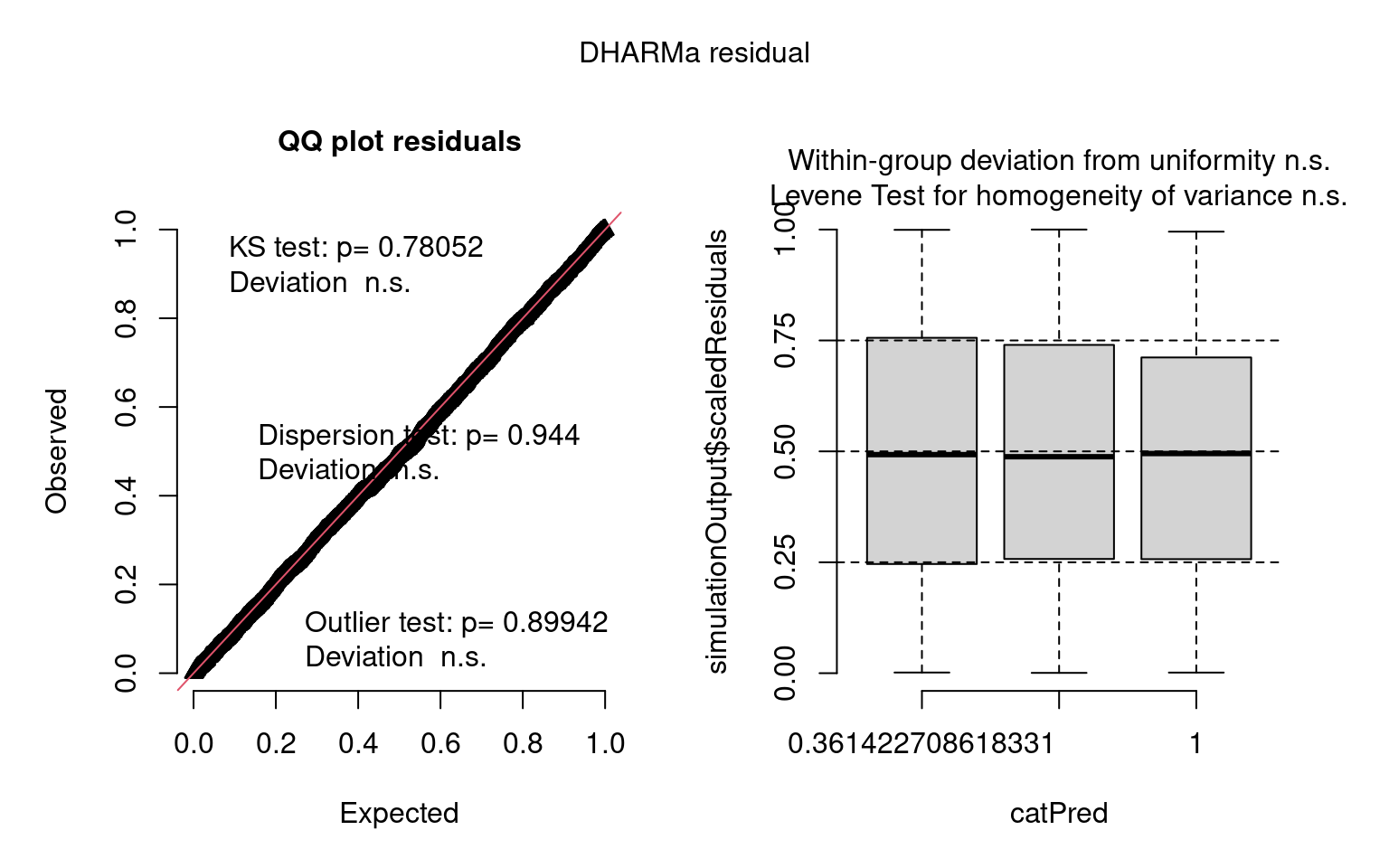

### unnamed-chunk-26-1.png

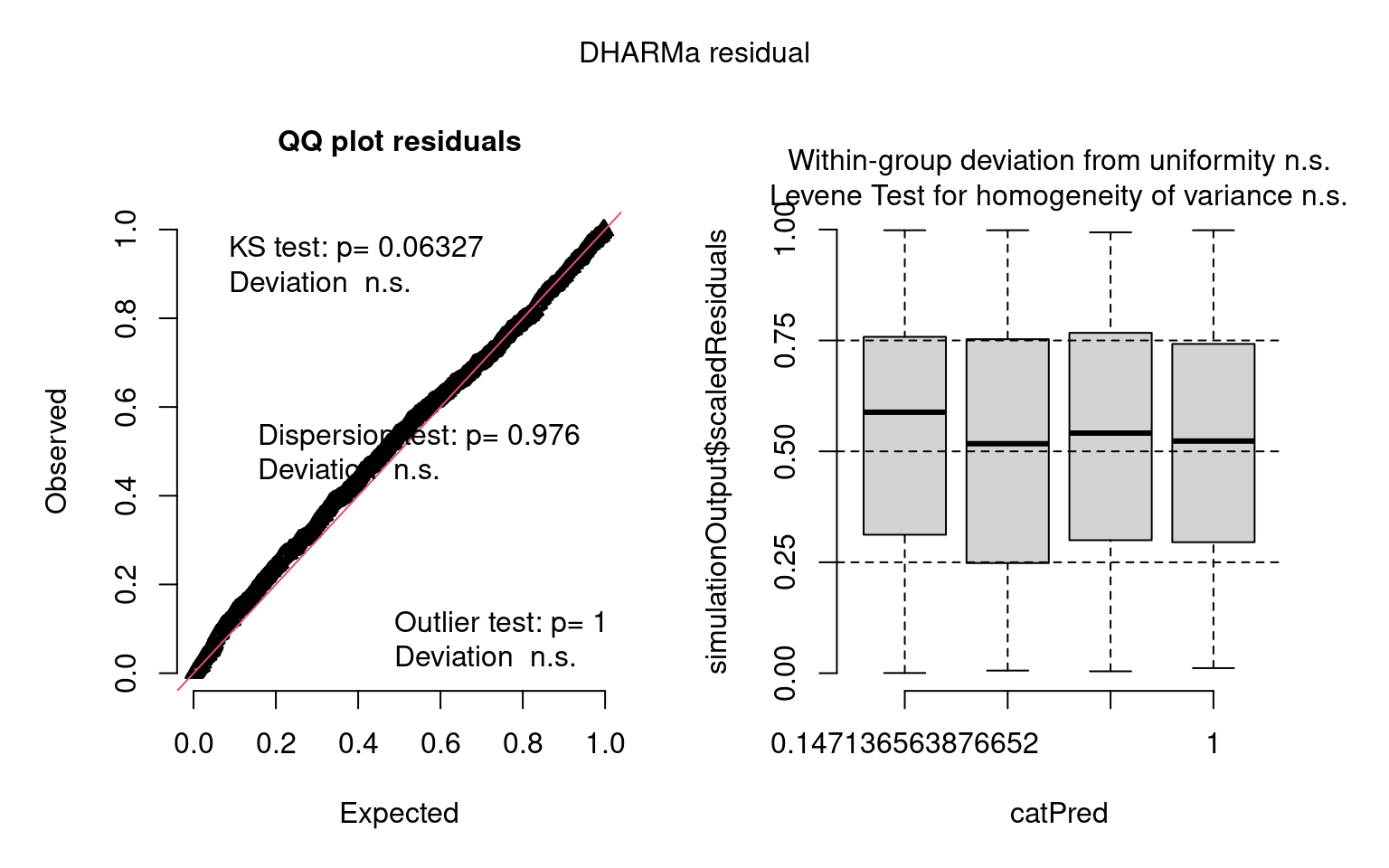

### unnamed-chunk-28-1.png

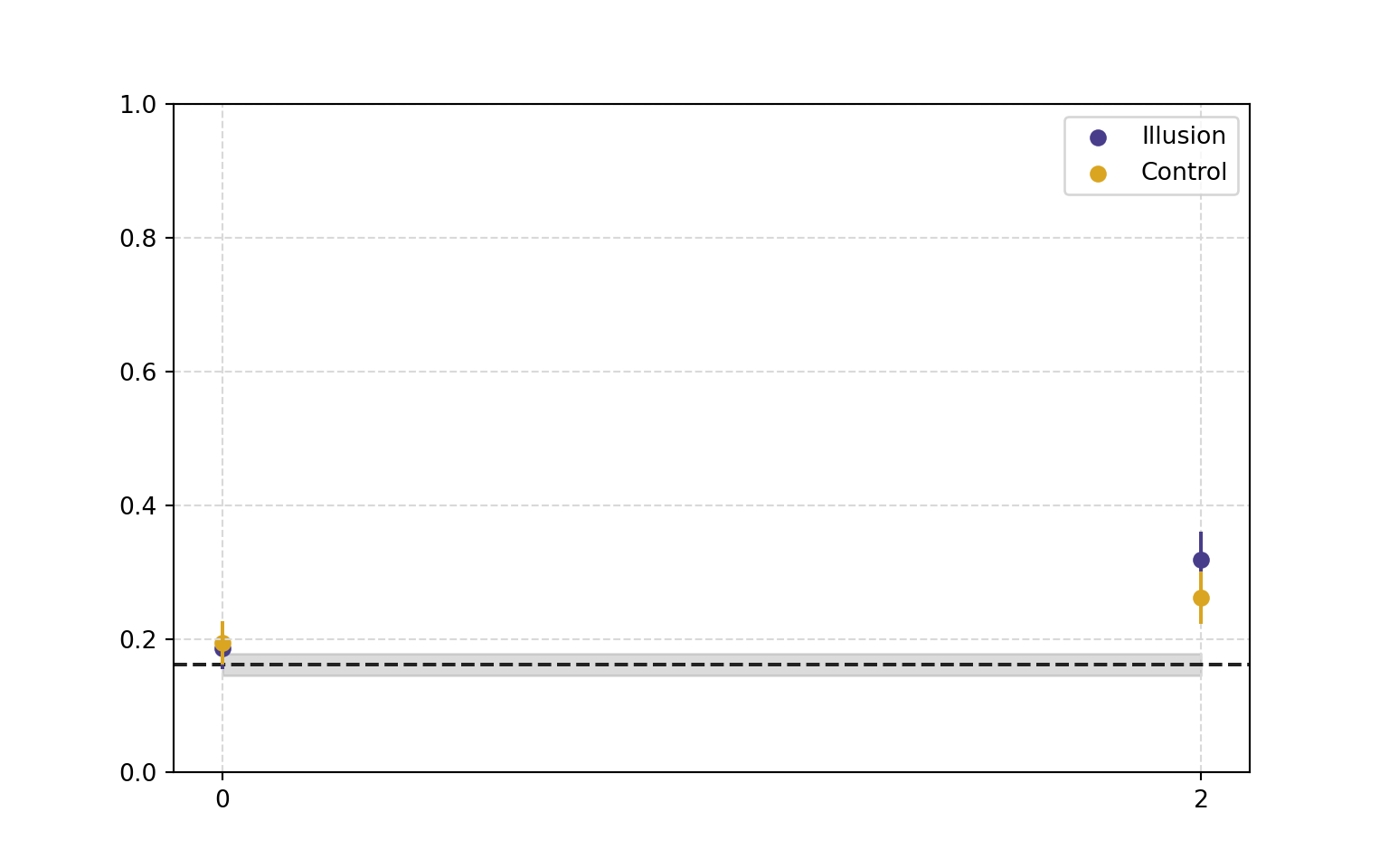

### unnamed-chunk-29-3.png

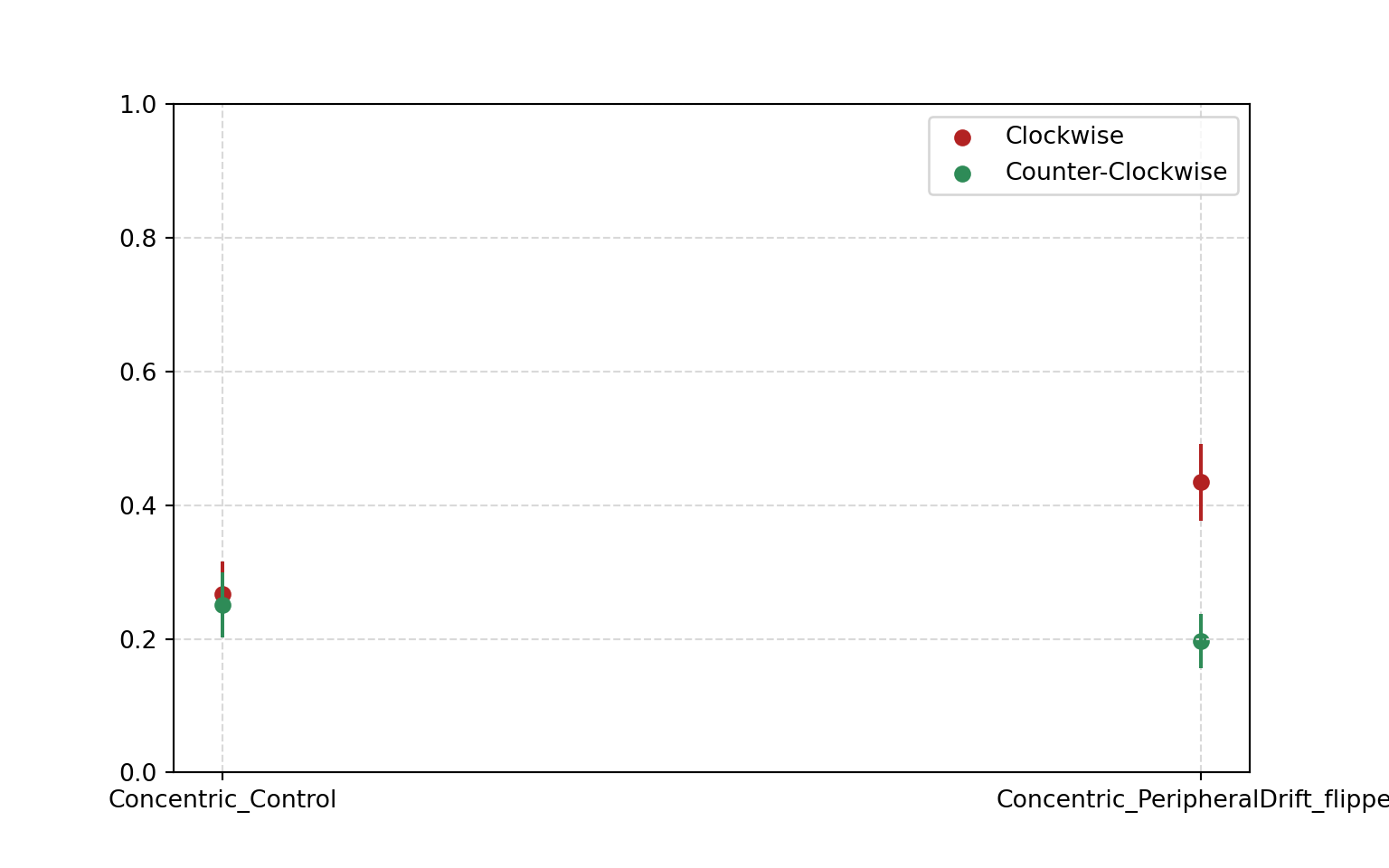
